# In Vivo Monitoring of Cellular Senescence by Photoacoustic and Fluorescence Imaging Utilizing a Nanostructured Organic Probe

**DOI:** 10.1101/2023.07.12.548691

**Authors:** Andrew G. Baker, Hui-Ling Ou, Muhamad Hartono, Andrea Bistrović Popov, Emma L. Brown, James Joseph, Monika Golinska, Chandan Sanghera, Estela González-Gualda, David Macias, Thomas R. Else, Heather F. Greer, Aude Vernet, Sarah E. Bohndiek, Ljiljana Fruk, Daniel Muñoz-Espín

## Abstract

Senescent cells accumulate in multiple age-related disorders, including cancer, exacerbating the pathological manifestations, and the eradication of these cells has emerged as a promising therapeutic strategy. Despite the impact of senescence in diseases, the development of tools to monitor the senescent burden *in vivo* remains a challenge due to their suboptimal specificity, translatability, and tissue penetrance. Here, we have designed a nanostructured organic probe (NanoJaggs) based on biocompatible indocyanine green dye (ICG) building blocks forming J-aggregates, which possess distinct spectral properties allowing both fluorescence and photoacoustic tomography (PAT) detection. We show that NanoJaggs are taken up by an active process of endocytosis and exhibit selective accumulation at the lysosomal compartment in several *in vitro* models for senescence. Finally, NanoJagg probe is validated in two *in vivo* studies including live PAT imaging and shows remarkable specificity to tumours with chemotherapy-induced senescence compared to untreated proliferative tumors. *In vitro, ex vivo* and *in vivo* all indicate that NanoJaggs are a clinically translatable tool for detection of senescence and their robust PAT signal makes them suitable for longitudinal monitoring of the senescent burden in solid tumors after chemo or radiotherapy.

## INTRODUCTION

Cellular senescence is a response to irreparable damage and stress that results in stable cell cycle arrest to prevent the expansion of altered (potentially pathological) cells. This program triggers the secretion of a complex mixture of inflammatory and tissue remodeling factors (senescence-associated secretory phenotype, known as SASP) to aid the repair of the surrounding tissue ^1^. Senescence also operates in response to oncogenic stress and, together with apoptosis, it has been defined as a mechanism of tumor suppression ^2^. Even during embryonic development, senescence plays an active role in tissue remodeling and promotes morphogenesis and organogenesis ^3, 4^. A common feature of the aforementioned physiological, tissue reparative, regenerative and tumor suppression roles is the recruitment of inflammatory cells to drive a clearance process of senescent cells that is ultimately executed by phagocytic cells ^1, 5^. However, persistent damage and stress can result in the deregulation of immunosurveillance and the accumulation of senescent cells in tissues, where they can contribute to the onset and progression of multiple age-related and chronic disorders and their pathological manifestations including, but not restricted to, cardiovascular diseases, fibrosis, neurological disorders, type 1 and 2 diabetes, obesity, inflammatory syndromes, musculoskeletal conditions and also cancer, by a number of cell autonomous and non-cell autonomous activities ^6, 7^.

After decades of research, the significance of cellular senescence in pathology has been acknowledged by the inclusion of this process within the hallmarks of both ageing and cancer ^8–10^. Although senescence can be a context-dependent antagonistic response, concluding studies have shown that the removal of senescent cells by pharmacogenetic approaches can significantly promote both healthspan and lifespan (by ∼30%) in progeroid and naturally-aged mice ^11, 12^. These landmark studies opened new therapeutic strategies aimed at developing pharmacologically active compounds to either remove senescent cells (senolytics) or modulate the pro-inflammatory SASP (senomorphics or senostatics). For example, it has been reported that the combination of Dasatinib and quercetin can improve the lifespan by 30% and reduce the mortality hazard by ∼65% in naturally-aged mice and senescent cell-transplanted young mice models ^13^. An arsenal of senolytic and senomorphic agents, including strategies to modulate the immune system and clear senescent cells by the use of CAR-T cells ^14^, have been reported since then with over 20 clinical trials currently in different stages of completion ^2, 7, 15–18^. Remarkably, in the context of cancer, both pro-senescent and anti-senescent therapies (the so called one-two punch combination strategy) have been shown to result in beneficial outcomes ^2, 18–20^.

Considering the role and impact of cellular senescence in disease, and an emerging field of senotherapies, surprisingly little progress has been made in the design and validation of contrast agents and tools for *in vivo* detection of the senescent burden. Unfortunately, there is no universal biomarker of senescence and therefore current strategies for the assessment of senescent cells in tissues take on a multistep or algorithmic approach using a number of biomarkers ^21, 22^. These commonly include the expression of cell cycle inhibitors/tumor suppressors (such as p16, p21, or p53) to identify stable cell cycle arrest, increased lysosomal senescence-associated β-galactosidase (SA-β-gal) activity, as well as senescence-associated heterochromain foci to identify nuclear structural changes and the expression of SASP factors ^22, 23^. However, such a strategy is only suitable for *ex vivo* biopsied samples ^17^, which limits potential applications for real-time or longitudinal senescence monitoring in human settings.

A number of small molecule fluorescent probes incorporating switchable ‘turn on” systems were designed by utilizing the enhanced activity of lysosomal β-galactosidase ^24–26^. In addition, a distinct strategy based on targeting mitochondria using a membrane permeable cyanine dye (CyBC9) was explored to detect senescent mesenchymal stromal cells ^27^. However, fluorescence is limited by the penetration depth of the excitation and emission light (µm to mm) making it in many cases ineffective for potential clinical applications, in particular for imaging internal organs that require a penetration depth of up to several centimeters ^28, 29^. As an *in vivo* alternative to fluorescence, two positron emission tomography (PET) probes have been reported ^30, 31^, both of them based on the exploitation of the increased SA-β-gal activity in senescent cells. As a potential limitation, this strategy is based on radioactive tracers and sophisticated imaging systems. Finally, enhanced SA-β-gal activity is not a distinctive feature of senescent cells only, but also other cellular types like osteoclasts, neurons and macrophages ^32^.

To address the drawbacks of existing imaging strategies, photoacoustic tomography (PAT) has recently emerged as a powerful *in vivo* imaging technique ^33^, with significant potential for early detection of cancer ^34–37^. PAT employs a pulsed laser to excite the appropriate contrast agent, resulting in conversion of absorbed light into acoustic energy. Because acoustic waves scatter to a lesser extent than light, ultrasound images can be constructed from higher penetration depth compared to light microscopy techniques. PAT is able to optically map absorbance of either endogenous molecules such as hemoglobin, or contrast agents such as near-infrared (NIR) absorbing dyes ^38^. In contrast to computed tomography (CT) and PET, PAT does not require ionizing radiation, the acquisition times are shorter, and the instrumentation more affordable and easier to operate ^34^, albeit applied for localized imaging up to a few centimetres depth. Importantly, by acquiring images at several wavelengths, PAT allows for multiplexed imaging, resulting in data sets that reveal the distribution of several chromophores ^35^.

Applying PAT to the detection of senescent cells requires a photoacoustic contrast agent with a high molar extinction coefficient and a sharp, unique absorption spectrum to maximise the amount of light absorbed, and enable unmixing from other chromophores ^38^. In addition, the contrast agent should be photostable with a low quantum yield to ensure that the absorption spectrum does not change over time and most of the light energy is converted to heat ^39^. These requirements make nanoparticle (NP) systems particularly well-suited as they often have tuneable absorption spectra, high extinction coefficients and a large surface area that allows for further (bio)functionalization with stabilizing and targeting agents ^38^. The latter is important to enable the detection of specific cell types in highly heterogenous biological systems. However, in terms of senescent cells, despite many reports highlighting potential routes for their targeting ^40–47^, recent studies have sparked debate over the relative benefits of the use of targeted approaches *vs* non-targeted species ^48–51^.

Here, we have designed, synthesized and validated a novel nanostructured probe for *in vivo* detection of senescent cells, which is prepared by using FDA-approved indocyanine green (ICG) dye assembled into nanosized J-aggregates structures (NanoJaggs) under mild conditions. J-aggregates, initially described nearly 100 years ago, are supramolecular assemblies of organic dyes such as cyanines, porphyrins and perylene bisimides, characterized by unique optical and photophysical properties ^52, 53^. J-aggregates have been used in imaging (PAT, NIR and fluorescence microscopy) ^54, 55^, design of sensors ^56^, photonic devices ^57^ and as photothermal agents in cancer therapy ^54, 55^. Although their mechanisms of self-assembly and phenomena such as “superquenching” and “superradience” are still not fully understood ^52^, a number of nanostructured J-aggregates, ranging in sizes from 90 to 130 nm, have been prepared to date, mainly for tumor imaging and photothermal therapy ^55, 58–61^. Our study provides a simple strategy for the preparation of stable and pure J-aggregates that can be stored both in powdered form and in solution. In addition, we provide detailed physiochemical analysis of the composition of the probe and show that it can preferentially accumulate in lysosomes of senescent cells exploiting clathrin and macropinocytosis pathways of cellular uptake. Importantly, biocompatibility and exceptional contrast agent features are demonstrated in *in vitro*, *ex vivo* and *in vivo* by using tumor xenograft mouse models of chemotherapy-induced senescence exhibiting both fluorescence and PAT properties that make NanoJaggs suitable for potential translational applications in human settings.

## RESULTS

### Synthesis and characterization of NanoJagg probes

NanoJaggs were prepared by stirring an aqueous solution of ICG dye at 65 °C for 24 h (**Fig. 1a**). The reaction was monitored by measuring the light absorbance of J-aggregates at λ_max_ = 895 nm and ICG at λmax = 780 nm (**Fig. 1b**). Following the completion of the reaction, which was characterized by loss of ICG absorption peak, purification by dialysis and ultra-centrifugation resulted in precipitation of pure NPs (see **Materials and Methods** for details). Although similar protocols have been reported before ^55, 58–60^, our methodology includes a purification (centrifugation) that is essential to separate the NanoJagg NPs away from other side products **(Extended Data Fig. 1)**.

**Fig. 1.**
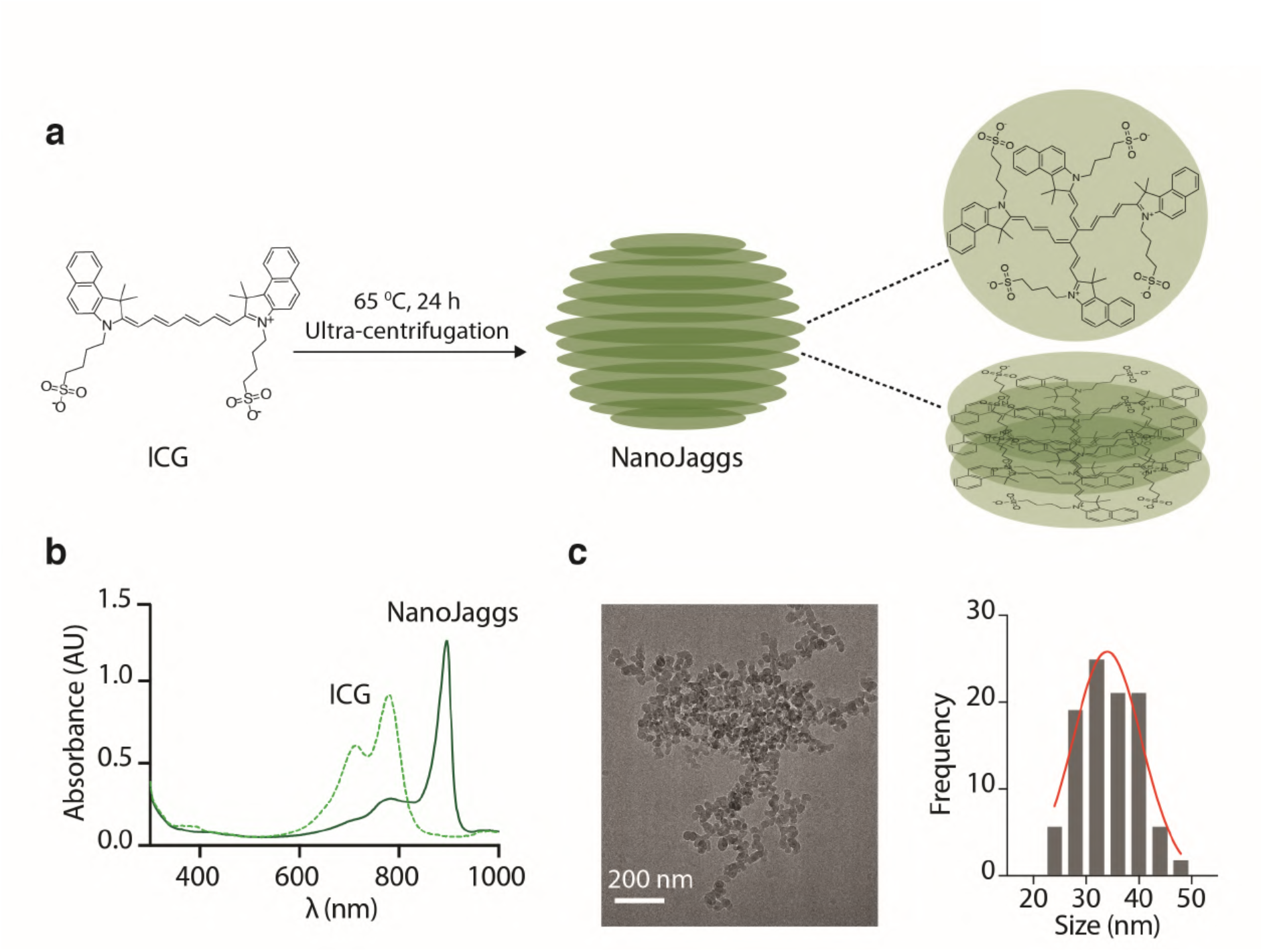
Synthesis and characterization of NanoJaggs. (**a**) Schematic overview for the preparation of NanoJaggs from ICG in an aqueous solution at 65°C for 24 h. The resulting product is purified using ultra-centrifugation (>15,000 rpm). (**b**) Characteristic red absorbance shift from 780 to 895 nm is observed upon J-aggregate formation (**c**) Cryo-TEM image of the NanoJagg NPs with an average diameter of 34±4.8 nm.

Both NMR and LC-MS/MS analysis was performed to gain better understanding of the structural composition of the pellet and supernatant (**Extended Data Fig. 2 to 4**). A single peak with 751 m/z was observed in the chromatogram of the pellet (**Extended Data Fig. 3b**), while the chromatogram of the supernatant contained multiple peaks, some of which have been identified as ICG degradation products. The disappearance of the ^1^H-NMR peak at 6.60 ppm in the NanoJaggs, which corresponds to the hydrogen in the carbonyl chain of ICG, indicates the formation of the bridging bond between two ICG molecules similarly to what was previously reported for an ICG dimer (**Extended Data Fig. 3**) ^62, 63^. On the other hand, the ^1^H NMR of the supernatant confirmed the presence of multiple compounds (**Extended Data Fig. 4**) and we also confirmed that the degradation products make up 54% of the total yield by mass. Previously reported J-aggregate nanostructures were used as prepared and not purified, suggesting that they were complex mixtures of ICG degradation products and J-aggregate NPs. Unlike those mixtures, our pellet was shown to be a chemically pure nanostructure composed of ICG dimers (**Extended Fig. 1b**). HPLC assessment of the dimer content showed that 97% of NanoJaggs are dimer structures, in contrast to the complex mixture of compounds found in the supernatant (**Extended Data Fig. 5**).

To gain a better understanding of the size distribution and shape of the obtained NanoJaggs, cryo-transmission electron microscopy (TEM) imaging was performed showing semi-spherical NPs with an average diameter of 34±4.8 nm (**Fig. 1c and Extended Data Fig. 6b**). Further, Energy dispersive X-ray analysis (EDX) demonstrated the presence of sulfur stemming from the sulfonate group on the ICG building blocks.

Spectrally, in addition to the red shift in absorbance of NanoJaggs compared to ICG (λ_max_ = 895 nm and λ_max_ = 780, respectively) seen in **Fig. 1b**, the fluorescence quenching of ICG dimers within the aggregates (λ_em_ = 810 nm) was also demonstrated. (**Extended Data Fig. 6c and d**). Such quenching is beneficial for photoacoustic imaging as it results in a larger percentage of absorbed energy being transformed into acoustic energy, thus enhancing the PAT signature^64^. In fact, when normalized to the same absorbance maximum, NanoJaggs show a three-fold enhancement in total photoacoustic signal generated (**Extended Data Fig. 6d**), which is in line with previous reports on photoacoustic properties of J-aggregates ^65^. In addition, the extinction coefficient was found to be 1.92±0.07 × 10 9M^-1^cm^-1^ (**Extended Data Fig. 6e**), which is similar to that of metallic plasmonic NPs ^38, 66^, despite NanoJaggs being entirely made of organic molecules. Finally, no changes were observed in absorbance spectra of NanoJaggs evaluated in various biologically relevant buffers (PBS at pH 7.4, DMEM, DMEM with 10% FBS and FBS) over 7 days indicating their remarkable colloidal stability (**Extended Data Fig. 7**). Altogether, these results demonstrate efficient and reproducible method to synthesize ultrapure, soluble and highly stable nanostructured NanoJaggs probes with fluorescent and PAT properties.

### NanoJaggs accumulate in the lysosomal compartment in senescent cells

Previous reports have shown that ICG is efficiently taken up into lysosomes of cancer cell lines, ^67^ and can accumulate in solid tumors *via* (often disputed) enhanced permeability and retention (EPR) effect ^67, 68^. Compared to cancer cells and normal (differentiated) cells, senescent cells are characterized by an enlarged lysosomal compartment reflected in both an increased size and number of lysosomes. As a consequence, enhanced activity of lysosomal SA-β-gal at pH 6.0 is used as one of the most important biomarkers of senescence ^5, 23, 69, 70^. Based on the accumulation profile of ICG, we hypothesized that ICG containing NanoJaggs may also be taken up and localized at lysosomes in cancer cell lines, and that this property would make NanoJaggs an efficient probe for detection of senescent cells, in particular therapy-induced senescent cancer cells characterized by high lysosome content. To test this hypothesis, cell uptake and colocalization studies by confocal microscopy with NanoJaggs were first performed in human A549 (lung adenocarcinoma) and SK-MEL-103 (melanoma) cancer cell lines showing that NanoJaggs, similarly to ICG, can target the lysosomal compartment (**Extended Data Fig. 8**).

We next sought to ascertain the selectivity of NanoJaggs for senescent cells. Since senescence can be a heterogenous response that depends on the trigger, cellular type and context, a variety of senescent cell models were employed, namely: Palbociclib-induced senescence (by selective inhibition of cycling dependent kinases 4 and 6, CDK4/6) in SK-MEL-103 and A549 cancer cells, Cisplatin induced senescence (by using a DNA intercalation chemotherapy) in A549 cancer cells, and radiation induced senescence (by DNA damage and double strand breaks induction) in human WI-38 fibroblasts (**Fig. 2a**). The implementation of a *bona fide* senescent status was assessed by standard protocols, including increased SA-β-gal activity, stable arrest of cell cycle progression by proliferation assays, and by enhanced expression of p21 cell cycle inhibitor and, conversely, the decreased expression levels of phosphorylated RB (a marker accounting for an active progression of the cell cycle) by Western blotting (**Extended Data Fig. 11**). Confocal microscopy experiments showed significantly increased accumulation levels of NanoJaggs across all senescent cell models when compared to their non senescent counterparts (**Fig. 2b and c**; **Movie 1 and Movie 2**). It should be noted senescent cells are characterized by an aberrant morphology and enlarged size in comparison to their own non-senescent cellular states^5^. Therefore, relative fluorescent units are normalized by the total cell area and hence the obtained average signal is comparable across different cell types. Largest NanoJagg uptake was observed in senescent SK-MEL-103 and A549cells (9.4x and 5.2x, respectively, when compared to their non-senescent cancer cell counterparts), while radiation-induced senescent WI-38 fibroblasts accumulated only 2.7x higher fluorescent levels than the corresponding non-irradiated cells (**Fig. 2C**).

**Fig. 2.**
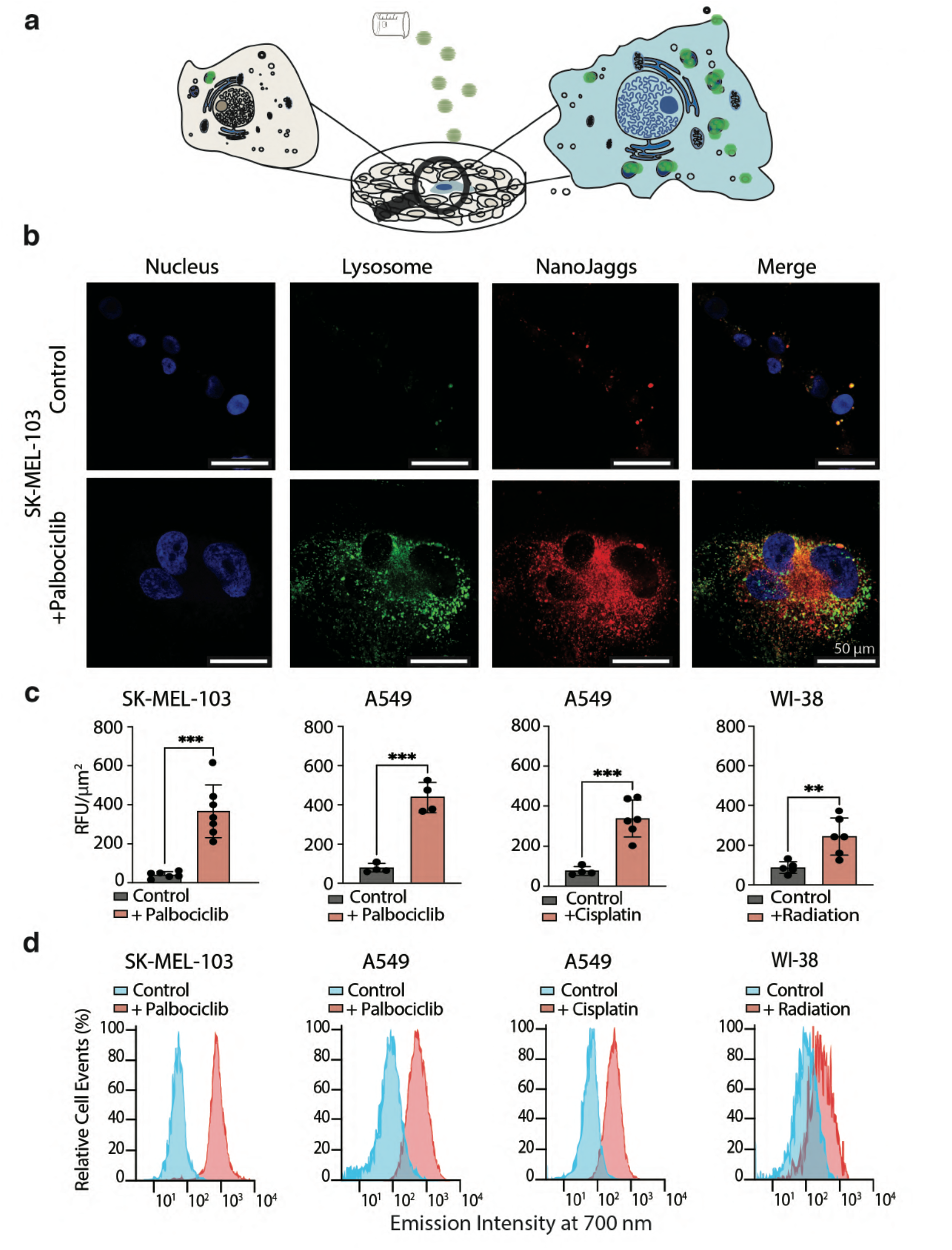
NanoJaggs are selective for senescent cells and accumulate in the lysosome. (**a**) Schematic representation showing the selective accumulation of NanoJaggs in senescent cells. (**b**) Confocal images of SK-MEL-103 non-senescent and senescent cells induced by Palbociclib and treated with 50 µg/mL NanoJaggs. (**c**) Confocal quantification of RFU/µm^2^ of the NanoJagg signal in the different cell lines. (**d**) Accumulation of NanoJaggs in cells evaluated with flow cytometry demonstrates increase in accumulation of senescent cells compared to controls. X-axis is log scale. Data represent as mean±SD, and a Two tailed t-test was used to calculate the significance (*p < .05, **p < .01, ***p < .001). All examples used show in significant increase in NanoJagg uptake compared to control cells and adjusted for cell area.

Accumulation of NanoJaggs in senescent cells, in contrast to cancer cells, was found to be dose dependent and exhibited a high level of sensitivity at the lowest concentration tested (0.4 μg/mL *vs* 50 μg/mL) (**Extended Data Fig. 9**). These results were further confirmed by flow cytometry studies showing a similar trend of NanoJaggs accumulation in Palbociclib-induced senescent SK-MEL-103 melanoma cells (12.5x higher relative levels of fluorescence emission intensity in senescent cells *vs* control cells), Palbociclib- and Cisplatin-induced senescence in A549 lung cancer cells (6x and 6.5x higher levels of fluorescence in senescent *vs* control cells, respectively) and in radiation-treated WI-38 senescent fibroblasts (2.5x higher levels of fluorescence in senescent *versus* control cells) (**Fig. 2d**).

Colocalization studies showed that NanoJaggs largely merged with lysosomes at the perinuclear space (**Fig. 2b and Extended Data Fig. 10 and 12**) whereas, in sharp contrast, there was no significant colocalization with mitochondria (**Extended Data Fig. 13**) or endoplasmic reticulum (**Extended Data Fig. 14**). Interestingly, selective accumulation of NanoJaggs in senescent cells could also be observed via conventional optical/light microscopy in both A549, and SK-MEL-103 cell lines, thereby making this senescent probe (senoprobe) comparable to commercially used SA-β-galactosidase activity kits in terms of selectivity (**Extended Data Fig. 15**), but with the advantage of being suitable for live cells due to the low levels of toxicity of NanoJaggs. Biocompatibility of NanoJaggs was assessed by CellTiter-Blue viability assays where senescent and non-senescent human lung adenocarcinoma (A549) and melanoma (SK-MEL-103) cell cultures were supplemented with a wide range of NanoJagg concentrations (0.1 – 100 µg/mL) (**Extended Data Fig. 16**).

Collectively, these data demonstrate that NanoJaggs are a non-toxic and versatile senoprobe to trace multiple senescent phenotypes by optical and fluorescence microscopy displaying a lysosomal localization.

### NanoJaggs are taken up by an active mechanism of endocytosis in senescent cells

Following the significant increase in NanoJagg accumulation and fluorescent/optical signal observed in senescent cells we sought to gain insights into the underlying mechanism of cellular uptake. Active endocytosis pathways can be subdivided into different categories, namely macropinocytosis (or phagocytosis), pinocytosis, receptor-mediated endocytosis (also known as clathrin-mediated endocytosis), and receptor-independent endocytosis (involving cholesterol-binding protein caveolin plasma membrane buds)^71^. Both pre- and post-uptake inhibitors were employed to block these pathways and discern the possible routes by which NanoJaggs accumulate in lysosomes of treated senescent cells (**Fig. 3a**). Pre-uptake endocytosis inhibitors (Pitstop 2, Dyngo4a and phophatidylinostitol-3-kinase inhibitor Ly294002) either block cell membrane-based pathways such as macropinocytosis (Ly294002) and clathrin-coated pit formation (Pitstop 2) or inhibit the formation of dynamin protein (Dyngo4a) involved in clathrin- and caveolin mediated endocytosis ^72–75^ (structures in **Extended Data Fig. 17**). Post-uptake inhibitors interfere with processes within the cell such as later stage of clathrin-coated vesicle fission from the membrane (PMZ inhibitor) ^72^ and autophagy (Chloroquine) ^76^.

**Fig. 3.**
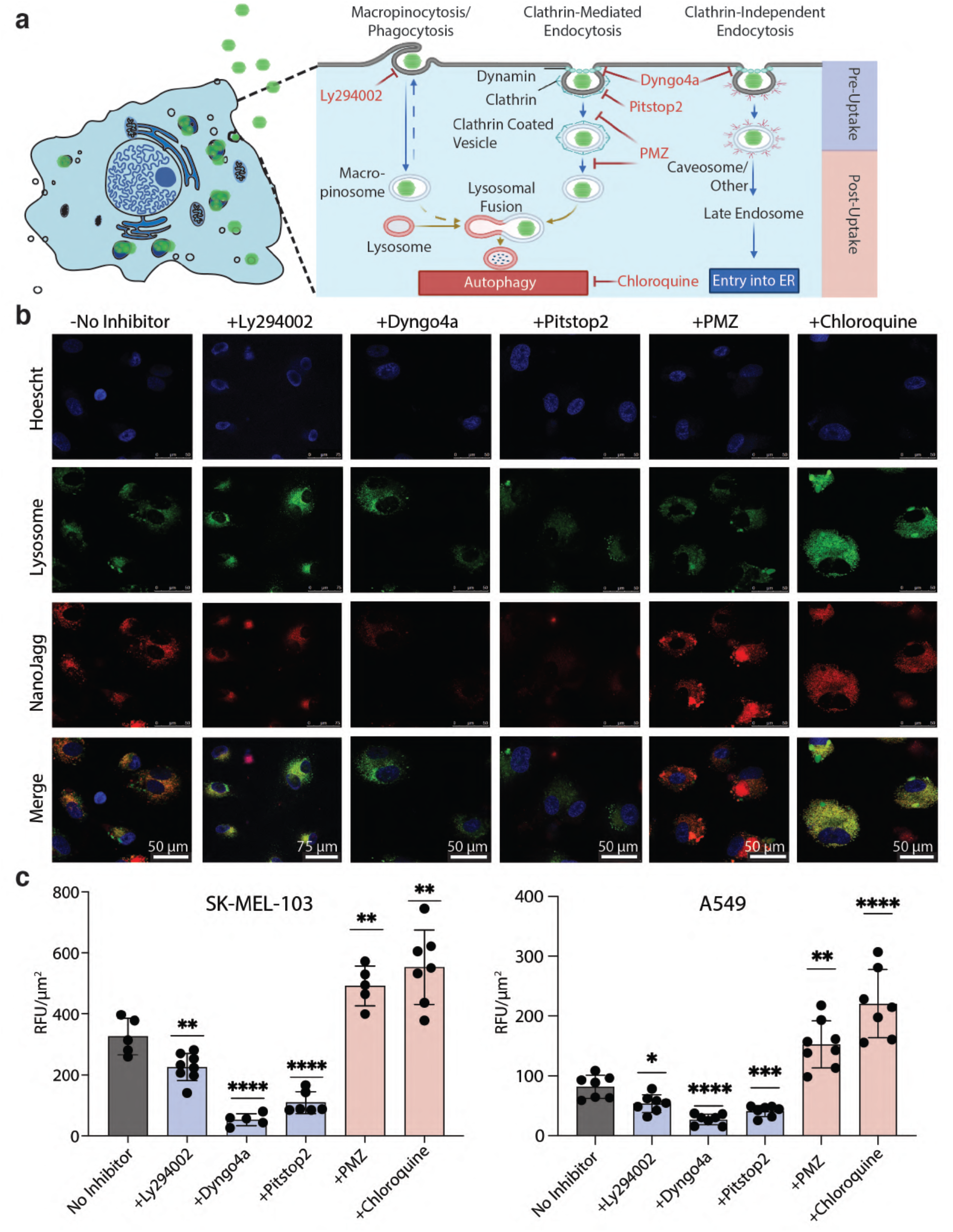
NanoJaggs target clathrin and micropinocytosis routes and remain in lysosomes. (**a**) Schematic representation of endocytosis in senescent cells and the inhibitors used in this study. Three pre-uptake inhibitors were used (Pitstop 2, Dyngo4a and Ly294002), as well as two post-uptake inhibitors (PMZ and Chloroquine). (**b**) Confocal images of NanoJagg uptake in senescent SK-MEL 103 cells treated with pre- and post-uptake inhibitors of endocytosis. Lysosomes are stained with LystoTracker green and nuclei with Hoechst (blue). (**c**) Confocal quantifications (RFU/µm^2^) of SK-MEL-103 and A549 cells. All comparisons are made to cells not treated with inhibitors, shown in black. Pitstop 2 and Dyngo4a both result in a significant inhibition of NanoJagg uptake in both cell lines (p < 0.0001), while Ly294002 caused a less but still significant reduction (p < 0.01 for SK-MEL-103 and p < 0.05 for A549). Both post-uptake inhibitors cause an increase in the NanoJagg signal: prochlorperazine’s increase p < 0.01 for SK-MEL-103, and p < 0.05 for A549; Chloroquine causes p < 0.01 for SK-MEL-103 and p < 0.0001 for A549. Data represent mean ± SD and a Two-tailed t-test was used to calculate the significance (*p < .05, **p < .01, ***p < .001, and ****p<0.0001).

Confocal microscopy studies showed significant reduction in NanoJagg uptake (p<0.0001) by senescent SK-Mel-103 melanoma and A549 lung cancer cells in presence of inhibitors of clathrin mediated endocytosis (Pitstop 2 and Dyngo4a) (**Fig. 3b and c, Extended Data Fig. 18 and 19**). In addition, macropinocytosis inhibitor Ly294002 also showed an effect on the NanoJagg uptake by senescent cells (p<0.01), although to a much lesser extent than the inhibitors of clathrin-mediated endocytosis (**Fig. 3b and c, Extended Data Fig. 18 and 19**). However, unlike senescent cancer cells, control (non-senescent) SK-MEL-103 and A549 cells treated with pre-uptake endocytosis inhibitors showed no significant differences in NanoJagg intracellular accumulation (**Extended Data Fig. 20 and 21**) indicating that the observed increased NanoJagg endocytic uptake is a senescence-dependent effect. Conversely to pre-uptake endocytosis inhibitors, both downstream inhibitors PMZ and Chloroquine, resulted in increased NanoJagg fluorescence levels in senescent SK-Mel-103 melanoma and A549 lung cancer cells (**Fig. 3b and c, Extended Data Fig. 18 and 19**), which were also enhanced in their non-senescent counterparts (**Extended Data Fig. 20 and 21**), thereby indicating that this is a senescence-independent effect. Increased levels of NanoJaggs fluorescence and maintenance of lysosomal accumulation upon Chloroquine treatment (**Extended Data Fig. 18, 19, 20, and 21**) are consistent with its reported effects resulting in lysosome dilation, impairment of autophagosome fusion with lysosomes and inhibition of autophagic flux ^77^.

Together, our results indicate that the NanoJaggs increased lysosomal accumulation in senescent cancer cells is, at least in part, mediated by an active process of endocytosis involving routes of macropinocytosis and clathrin-mediated endocytosis.

### NanoJaggs possess selective fluorescent properties in therapy-induced senescent xenografts *ex vivo*

To evaluate the efficiency of NanoJaggs to target senescent cells in a more biologically-relevant model we employed immunocompromised mice bearing tumor xenografts generated with SK-MEL-103 melanoma cells which were treated with senescence-inducing chemotherapy, as shown before^19^. Upon tumor formation, mice were treated daily with Palbociclib for 7 days (**Fig. 4a**), and this resulted in high levels of intratumoral senescence, as we inferred from elevated SA-β-gal activity (**Fig. 4b**), absence of the proliferative marker Ki-67, and reduction in phosphorylated Rb ^19^. Palbociclib-treated mice carrying SK-MEL-103 xenografts were given a single injection of vehicle (DMEM phenol red free), NanoJaggs (200 μL at 1 mg/mL) or the same amount of ICG dye and fluorescent levels were analyzed 6 h later by fluorescence imaging (**Fig. 4a**).

**Fig. 4.**
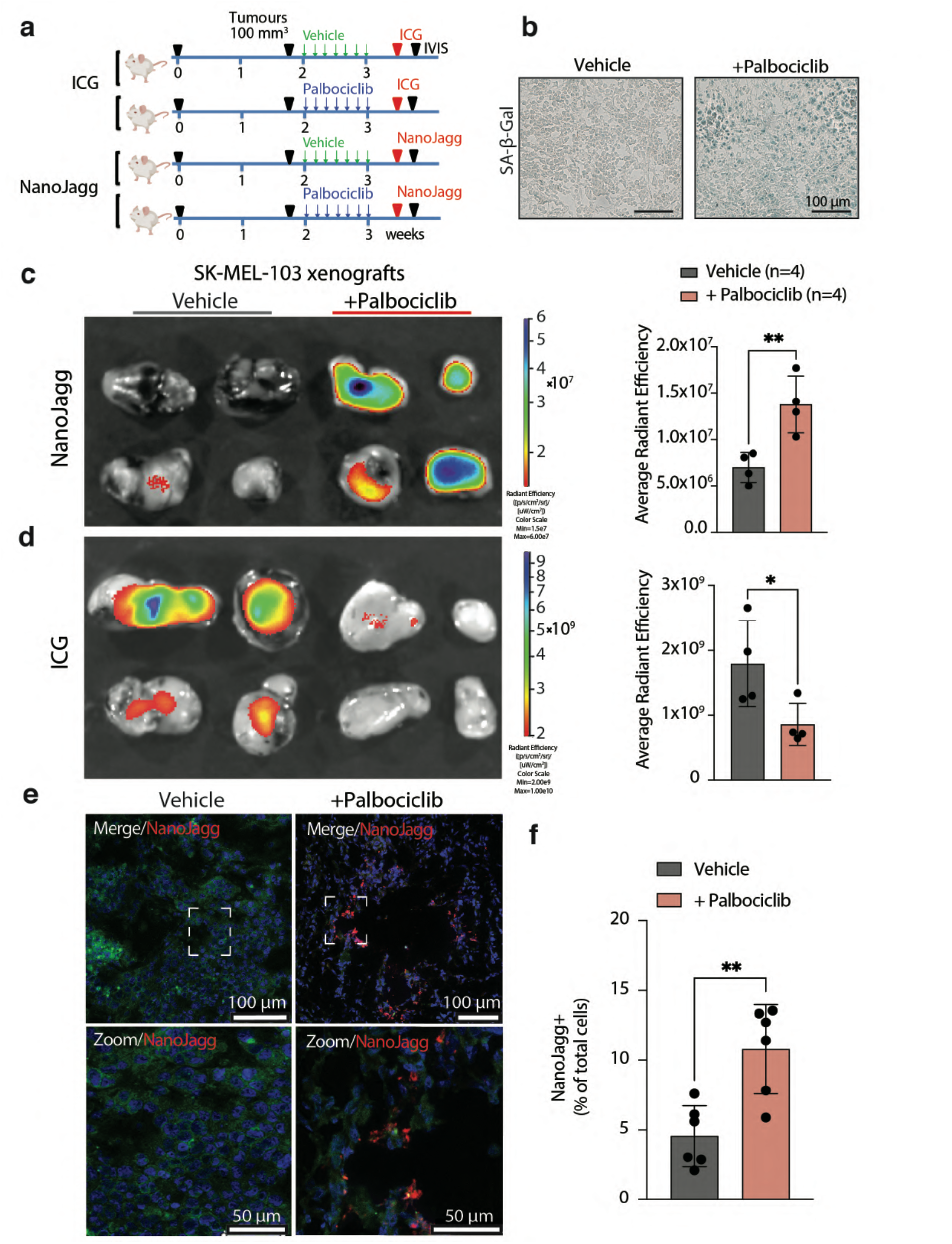
NanoJaggs target senescent tumors and ICG targets dividing tumors. **(a)** Schematic overview of the *in vivo* experiment. Briefly, the uptake of both ICG and the NanoJaggs were compared in Palbociclib-treated tumors *vs* untreated tumors. (**b**) Senescence associated β-galactosidase (SA-β- gal) staining of untreated and Palbociclib treated tumors. (**c**) IVIS images of dissected control and senescent tumors after treatment with NanoJaggs for 6 h. Quantification of the average radiant efficiency, comparing NanoJagg fluorescent signal (p = 0.0077). (**d**) IVIS images of dissected control and senescent tumors after treatment with ICG for 6 h. Quantification of the average radiant efficiency, comparing ICG fluorescent signal (p = 0.0438). (**e**) Confocal images and quantification of NanoJagg positive cells comparing the untreated to Palbociclib treated tumors. Data represent mean ± SD, and a Two-tailed t-test was used to calculate the significance. (*p < 0.05, **p < 0.01, ***p < 0.001).

Importantly, significant levels of NanoJaggs fluorescence were detectable by IVIS imaging in Palbociclib-treated (senescent) tumors (n=4) but not in control (proliferative) tumors (n=4) (p=0.0077) (**Fig. 4c**). In contrast, ICG fluorescence was predominantly observed (p=0.0438) in non-senescent tumors (n=4) compared to senescent tumors (n=4) (**Fig. 4d**), which is in agreement with previously reported EPR properties of ICG dye to accumulate in aggressive solid tumors (see *Introduction*). As an internal control, the background fluorescence signal of Palbociclib-treated (senescent) tumors in the absence of NanoJaggs was negligible at the tested conditions (**Extended Data Fig. 23a**), demonstrating that fluorescence can be predominantly attributed to the accumulation of NanoJaggs and ICG dye within the tumors. *Ex vivo* biodistribution studies showed that NanoJaggs fluorescence was hardly detectable after 6 h post-administration in organs of the reticuloendothelial system, including the liver and spleen, in untreated and Palbociclib-treated mice. In both groups, a considerable fluorescence signal was observed in the kidneys, suggesting that renal clearance can be a possible and fast route of elimination (**Extended Data Fig. 21b**). To further verify the accumulation of NanoJaggs in the senescent tumors, confocal microscopy imaging on 10 μm tissue slices was conducted confirming the preferential accumulation of the NanoJaggs in Palbociclib-treated (senescent tumors) (**Fig. 4e and f**, p=0.0028).

These results demonstrate that NanoJaggs, in contrast to ICG dye, can accumulate in tumors containing senescent areas in preclinical models and are detectable by displaying fluorescent properties.

### NanoJaggs exhibit potent PAT properties to detect a senescent burden *in vivo*

Next, NanoJaggs were explored as potential contrast agents for photoacoustic tomography (PAT) imaging of the senescent burden in therapy-treated tumors by using alive mice. PAT was applied to extend the potential imaging depth of accessible tumors while maintaining spatial and temporal resolution. To test NanoJaggs in PAT, we employed immunocompromised female nude mice bearing SK-MEL-103 melanoma tumor xenografts, either Palbociclib- or vehicle-treated, that were subsequently tail vein injected with NanoJaggs or ICG, and PAT imaging was performed at 6 and 24 h after injection (**Fig. 5a and b**). To validate the induction of senescence of Palbociclib-treated tumors, SA-β-gal activity assays and immunohistochemistry (IHC) staining of pRB and Ki-67 were conducted on dissected tumors after *in vivo* imaging. As expected, the Palbociclib-treated group showed an increased lysosomal senescence-associated β-galactosidase activity and a decreased nuclear staining of pRB and Ki-67, indicating a reduced proliferative capacity and the implementation of the senescent program (**Fig. 5c**). The loss of Ki-67 apparent in senescent tumors was further verified with immunofluorescence (**Extended Data Fig. 24a**). In addition, tumors treated with Palbociclib showed a marked reduction in proliferation rates, thereby confirming the onset of a stable cell cycle arrest (**Extended Data Fig. 24b to d**).

**Fig. 5.**
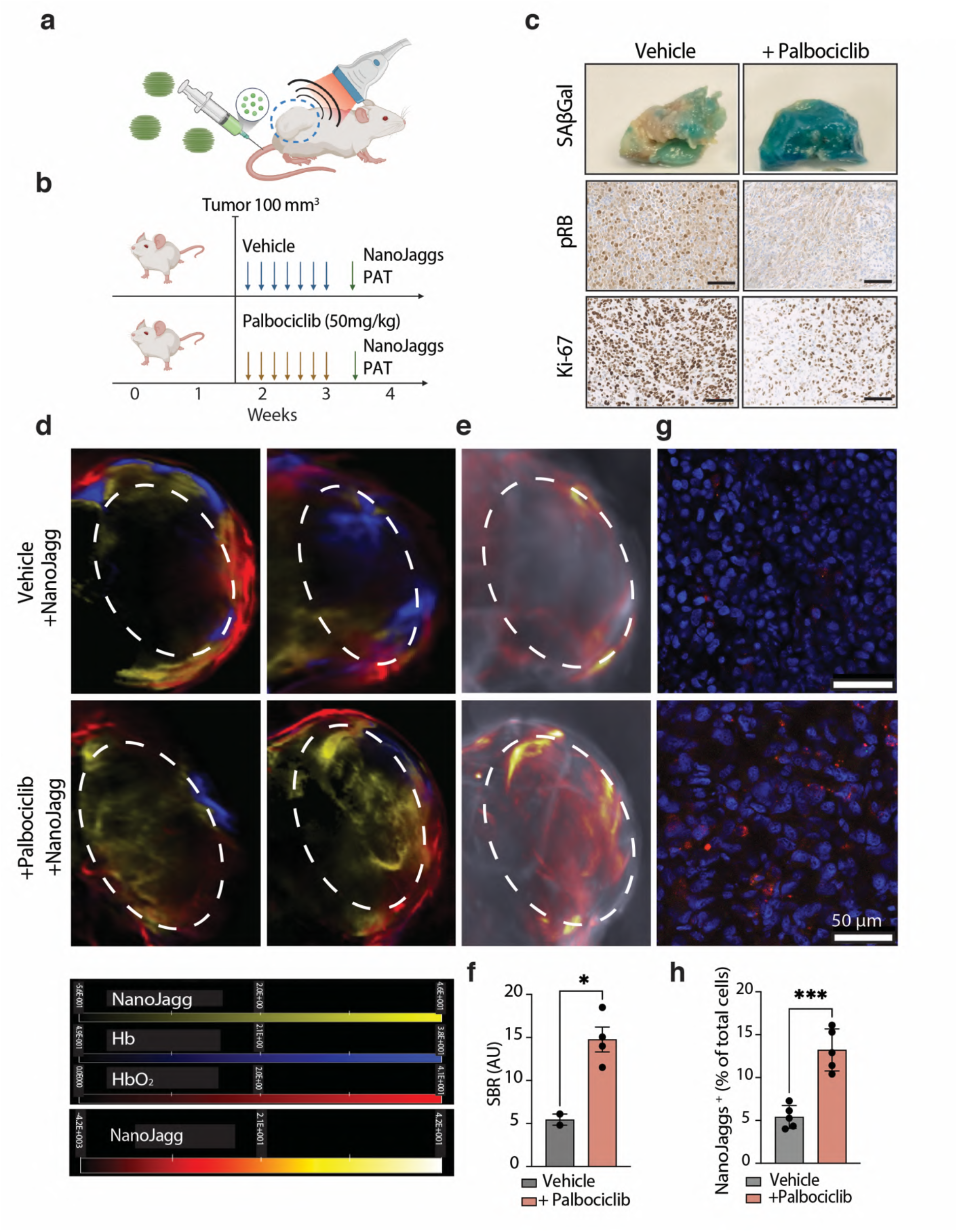
NanoJaggs act as contrast agent for *in vivo* for photoacoustic imaging of cellular senescence. (**a**) Illustrative cartoon of photoacoustic experiments. (**b**) Experimental design using two groups of mice treated and un-treated with Palbociclib both groups were given NanoJaggs after their treatment regime. (**c**) Tumors stained with X-gal show strong β-galactosidase activity in Palbociclib treated tumors. IHC images of Palbociclib treated tumor slices show a decrease in nuclear pRB and Ki-67 compared to control tumors. Scale bars 100 mm. (**d**) Images showing tumors spectrally unmixed for NanoJagg (yellow), Hb (blue) and HbO_2_ (red) scale bars shown below. (**e**) Representative axial images of control (n = 2) and senescent tumors (n = 4) treated with NanoJaggs scale bar shown below. Photoacoustic signal is in arbitrary units. SBR is signal/ background. (**f**) Quantification of NanoJagg SBR. A significant difference from the untreated *vs* Palbociclib treated tumors can be noted. (**g**) Confocal images and (**h**) quantification of NanoJagg positive cells comparing senescent and control mice. In both cases multiple tumors were used, as well as imaged in different parts of each slice. Data represent mean ± SD, and a Two-tailed t-test was used to calculate the significance (*p < .05, **p < .01, ***p < .001).

Linear spectral unmixing was applied to into hemoglobin (Hb), deoxyhemoglobin (HbO_2_), and NanoJagg/ICG spectra (**Extended Data Fig. 25a**). The potential of PAT imaging to be used for study of multiple data streams was demonstrated by analysis of de-oxygenated (Hb) and oxygenated (HbO2) hemoglobin distribution in and around tumor site. As it can be observed in **Fig. 5d**, NanoJaggs in non senescent tumors colocalize with Hb and HbO2 indicating vascular distribution. However, in the Palbociclib-treated (senescent) xenografts, NanoJaggs infiltrate the tumor and are localized away from the vasculature represented by Hb and HbO2 signals.

Signal to background ratio was determined from the signal of the tumor region of interest (ROI) divided by the signal of the background produced from spectral unmixing of the NanoJagg PAT spectrum. Representative axial images of control *vs* senescent tumors and the quantification of the NanoJagg photoacoustic signals are shown in **Fig. 5e, and f**. The signal to background ratio (SBR) was calculated at each axial plane of control and Palbociclib-treated tumors to represent the NanoJagg signal within the total tumor volume. Of note, a significantly higher SBR was observed in the senescent, Palbociclib-treated tumors (n=4), when compared to the proliferative tumor counterparts(n=2) (p=0.0135). This observation was further confirmed by the similar values obtained from the quantification based on contrast-to-noise ratios (**Extended Data Fig. 25b**). This analysis was repeated using both oxygenated (HbO2) hemoglobin and de-oxygenated hemoglobin (Hb) signals. In both cases no significant difference between senescent and non-senescent tumors (p=0.5099 and p=0.2926 respectively) was observed indicating that the NanoJagg signal accumulation is not due solely to increased blood perfusion of the tumors (**Extended Data Fig. 25c**). To further confirm that the NanoJagg signal is associated with senescence, a correlation analysis was conducted using both the tumor size and growth rate (**Extended Data Fig. 25d**). Remarkably, we found that the photoacoustic NanoJagg signal strongly correlates with the arrested tumor growth rate (R^2^ = 0.82, p = 0.002), but not the tumor size (R^2^ = 0.02, p = 0.794). This correlation was not observed in the case of endogenous hemoglobin HbO_2_, for which there was no correlation with growth rate (R^2^=0.069, p=0.6148, ns, **Extended Data Fig. 25e**).

To further confirm the presence of NanoJaggs in senescent tumors, confocal fluorescence microscopy images of the tumor slices were performed (**Fig. 5g, and h**). The total number of NanoJagg positive cells is significantly increased in senescent tumors compared to control counterparts (p=0.0002). In addition, these positive cells have significantly enhanced NanoJagg fluorescence signal than the untreated controls (p=0.0301, **Extended Data Fig 25f**).

Moreover, PAT was used to monitor biodistribution and routes of elimination of NanoJaggs confirming the data obtained from *ex vivo* studies (**Extended Data Fig. 26a**). PAT shows NanoJaggs accumulation in kidneys, suggesting renal clearance. However, contrary to the *ex vivo* images, NanoJagg accumulation can also be observed in the spleen and liver, possible due to the higher sensitivity of the technique. Importantly, NanoJaggs signal in the liver, spleen and kidneys decreases over time, and becomes significantly reduced after 24 h (**Extended Data Fig. 26a and b**). Finally, hematoxylin and eosin (H&E) staining showed no histological differences in the tissue architecture between vehicle- and NanoJagg-treated mice, showing no evidence of hepatic or renal damage and absence of spleen inflammation (**Extended Data Fig. 26c**).

Collectively, these results validate the use of a potentially translatable and biocompatible probe with high tissue penetrance exhibiting strong PAT properties for the detection of the senescent burden upon chemotherapy treatment in preclinical murine models.

## DISCUSSION

The biology of cellular senescence has gained much attention during the past decade due to the concluding evidence that this process contributes to the progression and even onset of many age-related disorders, including cancer, in multiple *in vivo* preclinical models^7^. Using the potential of nanostructured J-aggregates, we aimed to develop a probe selective for senescent cells that could be prepared in a cost-effective, reproducible, and scalable manner. To achieve this goal, several key challenges, that had not been addressed in previous studies, needed to be tackled. Unlike previously reported protocols ^55, 58, 60, 78^, which required several days for reaction completion, our strategy shortens reaction time to 24 h with further improvement possible at lower pH and different temperature (*data not shown*). Further, our synthetic strategy is highly reproducible and scalable, and delivers a nanostructured probe (NanoJaggs) stable in water and biologically-relevant media over several months.

In addition, NanoJaggs can be lyophilized and stored as a powder until they need to be used, significantly improving both the storage and transport. Most importantly, unlike previously reported systems, purification by ultracentrifugation and filtration resulted in NanoJaggs probes have a well defined and chemically pure structure. Both ^1^H NMR and LC-MS analyses showed that the main structural building block of NanoJaggs, with purity of 97% as determined by HPLC, are ICG-dimers.

While several degradation products of ICG can be found in the supernatant after ultracentrifugation of the reaction mixture, previous studies utilized J-aggregates without any purification steps. In fact, the reduced stability of these ICG J-aggregates in biological media such as PBS, DMEM, DMEM + 10% FBS and FBS compared to NanoJaggs probe described here, could be attributed to the presence of degradation products in employed non-purified systems. In the light of this information, previously reported protocols and probes, in particular those used in biomedical applications, should be re-evaluated and fully characterized. Spectroscopic studies of NanoJaggs provided more insight into the absorbance, fluorescence and photoacoustic properties of the probe, indicating fluorescence quenching and four times higher photoacoustic signal of NanoJaggs as compared to ICG, making this probe highly suitable for applications for *in vivo* photoacoustic imaging. Following the synthesis and characterization of NanoJaggs, which indicated high stability, and provided both fluorescent and photoacoustic signatures, we conducted *in vitro* uptake and co-localization studies by employing human melanoma (SK-MEL-103) and lung (A549) cancer cell lines, and human WI-38 fibroblasts, as well as their senescent counterparts (**Fig. 2**). Initial uptake studies using cancer cells showed that NanoJaggs accumulate preferentially in lysosomes (**Extended Data Fig 8**). Given that senescent cells have a significantly elevated lysosomal content and enzymatic activities, it was hypothesized that NanoJaggs would be ideal probes for senescent cells. To test this hypothesis, multiple therapy-induced senescent models were used, including Palbociclib-induced SK-MEL-103, Palbociclib-induced A549, Cisplatin-induced A549 and radiation-induced WI-38 cells. Both flow cytometry studies and confocal fluorescence imaging revealed a significant increase in NanoJaggs uptake in senescent cells when compared to their cancerous (non-senescent) counterparts, and the NanoJaggs fluorescent signal was found to substantially colocalize with the lysosomes but not with other organelles.

Cellular senescence can be a heterogenous – context dependent – response involving a variety of biomarkers. Recent studies have identified altered or aberrant endocytosis pathways as an emerging hallmark of senescence ^79^. To further elucidate the preferential uptake mechanism, senescent cells were treated with NanoJaggs in presence of various pre- and post-uptake inhibitors (**Fig. 3**). The uptake of NanoJaggs was significantly reduced by pre-uptake inhibitors such as, Pitstop2 and Dyngo4a, which affect clathrin-mediated endocytosis, suggesting that this pathway is the primary mechanism for their accumulation in senescent cells^73, 80, 81^. Additionally, Ly294002 data indicate that a measurable but probably smaller role is played by micropinocytosis ^74^. This aligns with previous studies that have shown an up-regulation of endo-lysosomal machinery as well as macropinocytosis (activated as a nutrient scavenging survival mechanism) and phagocytosis (used as a route of cell cannibalism) in senescent cells ^82–86^. In fact, senescent cells have often been linked to macrophages due to their ability to engulf other cells and fluorescent dextran ^82, 83, 87^, and this feature has been proposed to account for changes in the uptake machinery ^79^.

In contrast to pre-uptake inhibitors, the use of post-uptake inhibitors such as PMZ and Chloroquine resulted in an increased NanoJagg signal. However, there was no observed co-localization with lysosomes after PMZ treatment, which may be due to the interference of PMZ with the fusion of clathrin-coated vesicles and lysosomes^72^. In contrast, Chloroquine is an inhibitor of autophagy that results in lysosome dilation as it only interferes with processes downstream of lysosome formation and thus does not affect colocalization ^77, 88^. Together, these findings suggest that NanoJaggs are taken up by an active mechanism that likely involves clathrin-mediated endocytosis and macropinocytosis. This increase in uptake seems to be particularly pronounced in senescent cells, and although this phenomenon has been previously noted in the context of senescent cell biology, it has not yet been utilized for detection strategies or drug delivery.

With the goal to further evaluate the biocompatibility, efficiency to assess and monitor the senescent burden, and elimination routes of NanoJaggs, *in vivo* studies were conducted exploiting both their fluorescence and photoacoustic properties in murine models of cancer chemotherapy (**Fig. 4**). Using IVIS fluorescence imaging, preferential accumulation of NanoJaggs was observed in melanoma tumor xenografts undergoing Palbociclib-induced senescence when compared to vehicle-treated mice. Conversely, *ex vivo* analyses revealed accumulation of the ICG dye in (non-senescent) proliferating tumors, which is consistent with previous applications of ICG as a tumor contrast agent ^89, 90^. The presence of NanoJaggs signal in the kidneys 6 h after treatment suggests that renal clearance is a preferential route of elimination.

Building on promising findings from fluorescence imaging, the accumulation of NanoJaggs and their potential as PAT contrast agents was examined in a follow up study on live mice demonstrating a significantly increased PAT signal in senescent compared to the non-senescent tumors (**Fig. 5**). Furthermore, the use of PAT allowed for multiple data streams to be collected and analyzed, including blood oxygenation and blood vessel colocalization. The results showed that the NanoJaggs signal was distributed throughout the core of the tumors containing senescent areas, whereas in control (non senescent) proliferative tumors, NanoJaggs signal colocalized with Hb, indicating that the majority of NanoJaggs undergo circulation within blood vessels. PAT indicated that NanoJaggs can be eliminated through the reticuloendothelial system, including spleen and to a lesser extent, the liver, and are subjected to an efficient renal clearance. No signs of acute toxicity were observed after NanoJaggs treatment up to 24 h and the main organs showed no histological alterations. However, a more in-depth follow-up study on their biodistribution and maximum tolerated dose is planned to confirm these initial results.

These promising findings give an indication that NanoJaggs would have a route for visualization in a clinical setting. PAT is an emergent imaging modality offering significant translational applications in cancer and other disease^34, 37, 38, 91^. Two infrared (IR) dyes, ICG and cetuximab-800CW, are already being used as PAT agents in clinical studies for the mapping of lymph nodes in melanoma (NCT05467137) and cervical cancer (NCT03923881), respectively. In addition, several NP-based PAT contrast agents based on metal NPs and organic structures have been developed and validated in preclinical studies ^38^. Thus, imaging of NanoJaggs using PAT may provide a route to detect and characterize senescent cell populations *in vivo* should suitable biocompatibility and toxicity studies be undertaken to support clinical translation.

Our study has demonstrated that NanoJaggs can effectively serve as selective contrast agents for fluorescence and PAT imaging of senescent cells. Further research is needed to thoroughly understand the mechanism of NanoJaggs uptake and lysosomal targeting, but our preliminary data are consistent with previous findings showing altered endocytosis in senescence and hence support the view that active clathrin-mediated endocytosis pathways and macropinocytosis can play a role in the increased uptake of NanoJaggs by senescent cells. Also, since ICG contrast agents have been found to be preferentially taken up by cancer cells (in agreement with **Fig. 4D**) binding to the cell membrane followed by endocytosis ^67^), it is tempting to speculate that NanoJaggs might recognize specific factors of the senescent surfaceome allowing a selective targeting. The identification of the senescent surfaceome is an emergent field for the targeting of senescent cells that warrants further investigations. Importantly, with the recent advances in the development of drug nanocarriers and nanostructured contrast agents for healthcare applications, the demonstrated increase in NanoJaggs uptake by senescent cells could lead to the design of theranostics tools or selective nanostructured senolytics that do not require the addition of targeting species or functionalization strategies. Additionally, the targeting of NPs and senotherapies could potentially be enhanced or decreased using endocytosis inhibitors similar to those reported to enhance therapeutic antibodies in preclinical mouse cancer models and human trials_81_.

In conclusion, the NanoJaggs probe described here allows for the detection of multiple types of senescence through various techniques, including flow cytometry, confocal imaging, *in vivo* fluorescence and PAT imaging in different settings, including *in vitro*, *ex vivo*, and *in vivo* models. The enhanced accumulation of the probe in senescent cells correlates with their increased lysosomal content and we provide evidence that the preferential uptake is regulated, at least in part, by clathrin-mediated endocytosis and macropinocytosis. The simple synthesis of this chemically pure organic probe makes large-scale preparation and clinical translation suitable. However, further studies are needed, including a full biodistribution/biodegradation study in healthy mice and investigation of other models such as senescence-related chronic disorders, naturally-aged mice or models of oncogene-induced senescence. *In vivo* detection and monitoring of the senescent burden using NanoJaggs holds the promise, of not only giving insight into the level of tissue dysfunction, but also making a significant impact on the risk stratification and prognosis for patients in a variety of disorders, including cancer, and inform senolytic interventions. This new detection modality could aid differentiation of patients responding to therapy and those at risk of tumor relapse and metastasis. Importantly, the PAT properties of NanoJaggs and higher tissue penetrance represents a realistic opportunity to translate for the first time a senoprobe to human settings and early trials.

## MATERIALS AND METHODS

### Synthesis of NanoJagg probes

Aqueous ICG (Acros Organics, 10321541) solution (0.75 mM, 10 mL) was sonicated for 10 min and then heated to 65 °C under stirring (500 rpm). Formation of J-aggregates (λ_max_=895 nm) was monitored by UV-Vis spectrophotometer indicating the completion of reaction at 24 h, after which the reaction mixture was centrifuged and washed three times (17000 rpm/ 31000 g at 4oC for 30 min in a Sorvall LYNX 4000 high speed centrifuge). The pellet was redispersed in deionized water, filtered through a 0.2 µm filter, and lyophilized to obtain dark green NanoJagg NPs (5.4 mg, 54% yield). Lyophilization was carried out using a Telstar LyoQuest benchtop freeze dryer (0.008 mBar, −70 °C).

### NanoJagg characterization

UV-Vis absorption spectra were obtained with an Agilent Cary 300 Spectrophotometer, as well as, The Spark (TECAN), and Infinite 200 Pro (Tecan). Fluorescence emission spectra were obtained using a Varian Cary Eclipse Fluorescence Spectrophotometer as well as, The Spark (TECAN), and Infinite 200 Pro (Tecan). The hydrodynamic size and zeta potential of the NanoJagg NPs were measured using a Zetasizer Nano Range instrument (Malvern Panalytical).

To estimate the extinction coefficient for the concentration of NanoJagg NPs, we assumed spherical particles, partial overlap of the ICG dimer within the structure, and uniform internal molecule structure. We used the dimer of ICG’s molecular weight of 1502.56 Da and size was measured in PyMOL (Schrödinger) to first determine the number of dimers in the solution and then size. We then adjusted this concentration, based on the number of dimers per NanoJagg NP which we estimated to be ∼5,700 per NP.

To determine the chemical composition LC-MS and ^1^H NMR were conducted on the obtained NanoJaggs and the supernatant obtained during the centrifugation. LC-MS was performed using a Waters’ Xevo G2-S bench top QTOF and was performed by the Department of Chemistry Mass Spectrometry Service, University of Cambridge (UK). Samples were first dissolved in methanol to ensure the NanoJagg structure was dissolved. ^1^HNMR measurements were carried out using 400 MHz QNP Cryoprobe Spectrometer (Bruker) by the NMR service of the Department of Chemistry,

University of Cambridge. Samples were dissolved in deuterated methanol to make sure the NanoJagg structure was dissolved.

ICG: ^1^H NMR (400 MHz, MeOD): δ 8.23 (d, J = 8.6 Hz, 1H), 8.13 – 7.94 (m, 2H), 7.70 – 7.56 (m, 1H), 7.54 – 7.41 (m, 1H), 6.73 – 6.49 (m, 1H), 6.39 (d, J = 13.4 Hz, 1H), 4.25 (t, J = 6.5 Hz, 1H), 2.94 (t, J = 6.8 Hz, 1H), 2.25 – 1.74 (m, 6H). HRMS: calculated for C43H47N2O6S2 (M+H)^+^: Mass predicted:752.29; Found:752.2930 NanoJagg (pellet): ^1^H NMR (400 MHz, MeOD) δ 8.37 – 8.25 (m, J = 15.7, 8.0 Hz, 4H), 8.10 – 7.92 (m, 6H), 7.67 (dd, J = 18.5, 8.4 Hz, 4H), 7.56 – 7.43 (m, 4H), 6.53 – 6.41 (m, 2H), 5.91 (d, J = 13.9 Hz, 1H), 4.30 – 4.21 (m, 2H), 4.14 – 4.06 (m, J = 7.4 Hz, 2H), 4.02 – 3.92 (m, 2H), 2.91 – 2.75 (m, J = 14.8, 8.2 Hz, 5H), 2.14 (s, 5H), 2.07 (d, J = 8.2 Hz, 6H), 2.00 – 1.79 (m, 9H). HRMS: calculated for C86H92N4O12S42 (M+H)^+^: Mass predicted:751.28; Found:751.2880 Supernatant obtained from the reaction mixture after purification: ^1^H NMR (400 MHz, MeOD) δ 8.38 – 8.20 (m, 1H), 8.19 – 8.11 (m, 1H), 8.08 – 7.87 (m, 2H), 7.84 – 7.76 (m, 1H), 7.74 – 7.57 (m, 1H), 7.56 – 7.39 (m, 1H), 7.40 – 7.32 (m, 1H), 6.52 – 6.33 (m, 1H), 3.94 – 3.85 (m, 1H), 3.03 – 2.70 (m, J = 31.1, 24.2, 17.8, 10.9 Hz, 2H), 2.31 – 1.71 (m, 6H), 1.68 – 1.55 (m, J = 7.7 Hz, 1H), 1.38 – 1.08 (m, 1H).

### Cryo-TEM of NanoJagg probes

Cryo-TEM micrographs were obtained using a Thermo Scientific (FEI Company) Talos F200X G2 microscope operated at 200 kV. Images were recorded on a Ceta 4k x 4k CMOS camera and processed with Velox software. Specimens for investigation were prepared through vitrification by plunge freezing of the aqueous suspensions on copper grids (300 mesh) with lacey carbon film (Agar Scientific). Prior to use, the grids were glow discharged using a Quorum Technologies GloQube instrument at a current of 25 mA for 60 s. Suspensions of the samples (2.5 µL of a 1 mg/mL solution) were pipetted onto the grid, blotted using filter paper, and immediately frozen by plunging in liquid ethane utilizing a fully automated and environmentally controlled blotting device, Vitrobot Mark IV. The Vitrobot chamber was set to 4°C and 95% humidity. Samples after vitrification were kept under liquid nitrogen until they were inserted into a Gatan Elsa cryo holder and analyzed in the TEM at −178°C. Cryo-TEM was performed by Heather Greer of the Department of Chemistry Electron Microscopy Facility.

### Photoacoustic properties of NanoJaggs

Photoacoustic measurements were performed using a commercial PAT system (inVision256-TF; iThera Medical GmbH) and reconstructed using a linear model-based approach in their proprietary software. A tube of NanoJaggs was placed in the center of a tissue-mimicking phantom that mimics the optical and acoustic properties of biological tissues.^93^ A region of interest was drawn around the NanoJagg inclusion, and the mean photoacoustic signal was extracted at each wavelength. After matching λ_max_ UV-Vis for both to 1AU, the photoacoustic signals of the ICG monomer and J-aggregate were compared, using the following wavelengths: 700, 720, 740, 760, 770, 780, 790, 800, 820, 840, 860, 870, 880, 890, 895, 900, 905, 910, 920, and 940 nm.

### Control and senescent Cell lines

The A549 (human lung adenocarcinoma) cell line was obtained from the European Collection of Authenticated Cell Cultures (ECACC). The SK-MEL-103 (human melanoma) cancer cell line was acquired from the American Type Culture Collection (ATCC). These cell lines were maintained in DMEM (Sigma) and supplemented with 10% FBS. For senescence induction, SK-MEL-103 cells were supplemented with the same media containing Palbociclib (PD0332991, MCE.) at 5 µM for 7 days. A549 cells were supplemented with the same media containing 15µM cisplatin (Stratech) for 10 days or 10 µM Palbociclib (MCE, PD0332991) for 10 days. Wi-38 cells were purchased from American Type Culture Collection (ATCC). This cell line was maintained in MEM (Thermo Fisher) supplemented with 10% FBS, 2 mM l-Glutamine and 1 mM sodium pyruvate (Sigma). Wi-38 cells were induced with 10Gy of X-ray irradiation, then maintained for 10 days. All cell lines were incubated in 20% O_2_ and 5% CO_2_ at 37 °C. Cells were routinely tested for mycoplasma using the universal Mycoplasma Detection Kit (ATCC) or by RNA-capture ELISA. For experiments with cells, Cisplatin (Selleck Chemicals, S1166) was reconstituted in sterile PBS; Palbociclib (MCE, PD0332991) was reconstituted in DMSO.

### Flow cytometry

Flow cytometry was measured on an LSR Fortessa (BD - Becton Dickinson). For the uptake of NanoJaggs we used 6 well plates and for all cell lines used we seeded 200,000 cells/well. Once the cells were attached, culture medium was changed to DMEM supplemented with 0.2 % FBS and cells were incubated with the NanoJaggs for 12–16 h. The cells were trypsinized and resuspended in PBS buffer with 2% FBS. DAPI (Sigma Aldrich, D9542) 0.1 µg/mL was added to each sample to exclude dead cells. For each condition 10,000 live events were collected for each. The analysis of all flow cytometry data was performed using FlowJo v10 (Treestar, OR). Unstained control and senescent cells were used as reference for each independent experiment. For ICG the 640 nm laser was used with an emission window of 750–810 nm. For the NanoJaggs, the 640 nm laser was used with an emission window of 708–753 nm.

### Confocal microscopy

Cells were seeded in a 96 well plates m-clear plates (Greiner Bio-One #655087) 3,500 control cells/well and 5,000 senescent cells/well. After 24 h cells were incubated with NanoJaggs at concentrations of 10-100 µg/mL. Confocal images were acquired on a Leica SP5 confocal microscope using a 20X HCX PL APO 0.5 NA dry objective or a 40X HCX PL APO 1.3 NA oil immersion objective. Hoechst (ThermoFisher, H1399) was used to specifically dye the nucleus at 5 µg/mL. Lysotracker (Cell Signalling Technology, FM8783), Mitotracker (MitoTracker Green (Cell Signalling Technology, FM9074), and ERtracker (BODIPY™ FL Glibenclamide, Thermo Fisher, E34251) were detected by using excitation wavelength of 488 nm (Argon laser) and with a detection window between 510 and 530 nm. The organelle specific dyes were used according to their manuals. NanoJaggs were detected by using excitation wavelength of 633 nm (Argon laser) and with a detection window between 680– 720 nm. Cells without dyes and NanoJaggs were images as autofluorescence controls using the corresponding excitation and detection wavelengths. Images were analyzed with LAS AF Lite (Leica).

### Organelle colocalization analysis by confocal microscopy

Cells were trypsinized and seeded in a flat-bottom µ-clear 96-well plates (Greiner Bio-One, #655087) at a density of 3,500–5,000 control and 4,000–6,000 senescent cell/well. Once the cells were attached, culture medium was changed to DMEM supplemented with 0.2% FBS and incubated with NanoJaggs (100 µg/mL – 10 µg/mL) for 12–16 h. Afterward, cells were washed 3x with PBS for 5 min. Specific organelle stains LysoTracker green DND-26 (Cell Signalling Technology, FM8783), MitoTracker Green (Cell Signalling Technology, FM9074), ER-Tracker™ Green (BODIPY™ FL Glibenclamide, Thermo Fisher, E34251) were used according to the manufacturers protocol. For nuclei staining, Hoechst (ThermoFisher, H1399) at 5 µg/mL in PBS was added 10 min prior to analysis.

### Endocytosis analysis by confocal microscopy

Cells were trypsinized and replated in flat-bottom µ-clear 96-well plates (Greiner Bio-One, #655087). Cells were seeded at a density of 3,500-5,000 control and 4,000-6,000 senescent cells per well. Once cells were attached, culture medium was changed to DMEM without FBS and exposed to several different inhibitors for 1 h. Pitstop2 (10 µM, Abcam ab120687), Dyngo4a (10 µM abcam. ab120689), Chloroquine (50 µM, Tocris, 4109), Prochlorperazine (15 µM, Sigma, P9178) and LY2940020 (20–50 µM, Sigma, 440202). After incubation, NanoJaggs were added in DMEM supplemented with 0.2% FBS, in a range from of 10 to 100 µg/mL for 16 h. Afterwards cells were washed 3x with PBS for 5 min. Specific organelle stains lyso green DND-26 (Cell Signalling Technology, FM8783) was used according to their manuals. For nuclei staining, Hoescht (ThermoFisher, H1399) at 5.0 ug/mL in PBS was added 10 min prior to analysis.

### Mice Experiments

All mice were treated in strict accordance with the local ethical committee (University of Cambridge License Review Committee) and the UK Home Office guidelines. Tumor xenografts were established using SK-MEL-103, a melanoma cell line. Cells were trypsinized, counted with a hemocytometer, and injected subcutaneously (0.5×10^6^ cells in a volume of 100 µL per dorsolateral flank) in 8– to 10–week old athymic nude female mice (Hsd:Athymic Nude-Foxn1nu) purchased from Charles River. Tumor volume was measured every 2 days with a calipers and calculated as *V* = (& × (2)/2 where a is the longer and b is the shorter of two perpendicular diameters. Palbociclib (MCE, PD0332991). was dissolved in 50 mM sodium lactate at 10.0 mg/mL and administered by daily oral gavage at the indicated doses. 200 µL of NanoJaggs (1 mg/mL) and 200 µL of ICG (1 mg/mL) were injected via the tail vein in clear DMEM (ThermFisher, no phenol red).

### *Ex vivo* imaging by IVIS

An IVIS Spectrum Imaging System (Perkin Elmer Inc) was used for ex vivo fluorescence imaging. For *ex vivo* imaging, mice were sacrificed by cervical dislocation and organs and tumor xenografts were analyzed immediately after harvesting. ICG was detected using an excitation wavelength of 710–740 nm and emission bandpass from 810–830. NanoJaggs were detected using an excitation wavelength of 605 nm and emission of bandpass of 700–720 nm. Fluorescence imaging quantification was performed by Living Image 3.2 software (Perkin Elmer Inc). A region of interest area (ROI) was drawn over around the tumors. Fluorescence activity is measured as average Radiant Efficiency with the equation of [-/.//02/.1]/[µ4//02].

### *In vivo* Photoacoustic Imaging

Photoacoustic measurements were performed using a commercial PAT system (inVision256-TF; iThera Medical GmbH). Briefly, a tunable (660–1,300 nm) optical parametric oscillator, pumped by a nanosecond (ns) pulsed Nd:YAG laser, with 10 Hz repetition rate and up to 7 ns pulse duration is used for signal excitation. All mice were treated in strict accordance with the local ethical committee (University of Cambridge License Review Committee) and the UK Home Office guidelines. Tumor xenografts were established using SK-MEL-103 melanoma cells. Cells were trypsinized, counted with a haemocytometer, and injected subcutaneously (0.5*10^6 cells in a volume of 100 µL per dorsolateral flank) in 8– to 10–week–old athymic nude female mice (Hsd:Athymic Nude-Foxn1nu, Charles River). Tumor volume was measured every 2 days with a caliper and calculated as *V* = (& × (2)/2, where a is the longer and b is the shorter of two perpendicular diameters. Palbociclib (MCE, PD0332991) was dissolved in 50 mM sodium lactate at 10 mg/mL and administered by daily oral gavage at the indicated doses. 200 µL of NanoJaggs (1 mg/mL) and 200 µL of ICG (1 mg/mL) were injected by the tail vein in DMEM (Thermo Fisher, no phenol red).

Mice were anaesthetized using <3% isoflurane and placed in a custom animal holder (iThera Medical), wrapped in a thin polyethylene membrane, with ultrasound gel (Aquasonic Clear, Parker Labs). Mice were imaged using the wavelengths 700, 720, 740, 760, 780, 790, 800, 820, 840, 860, 870, 880, 890, 895, 900, 905, 910, 920, and 940 nm, with an average of 10 pulses per wavelength.

PAT data analysis was performed using ViewMSOT software (v3.6.0.119; iThera Medical GmbH). Model-based image reconstruction and linear spectral unmixing were applied on data in the 700–940 nm wavelength range to retrieve the relative signal contributions of oxy–(HbO2), deoxy–(Hb) hemoglobin, ICG and the NanoJaggs. Linear spectral unmixing was performed with published spectra for ICG, oxyhemoglobin, and deoxyhemoglobin as well as the NanoJagg absorption spectrum collected in this study. Regions of interest were drawn manually over the tumor area, and unmixing quantities were averaged over the entire tumor volume. A corresponding background region of interest was drawn near the back of the mouse for each anatomical plane.

Signal to background ratio was taken as the signal from the tumor ROI divided by the signal of the background ROI for each anatomical slice: 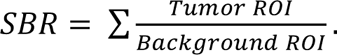 Contrast to noise ratio was calculated as the sum of the differences of the ROI signal and background divided by the standard deviation of the background signal for each tumor slice: 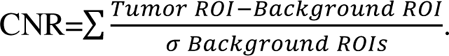

### Immunoblotting

Cell lysis was performed using RIPA buffer (Sigma) supplemented with phosphatase inhibitors (PhosSTOP™ EASYpak Phosphatase Inhibitors Cocktail, Roche) and protease inhibitors (cOmplete™ Protease Inhibitor Cocktail, Roche). Proteins were quantified and separated by SDS-PAGE and transferred to polyvinylidene difluoride (PVDF) membranes (Millipore) according to standard protocols. Membranes were immunoblotted with antibodies against p21 (556430) from BD Pharmingen, phospho-Rb (pRBS807/822) from Cell Signalling and normalized to GAPDH (ab9485) from Abcam. After incubation with the primary antibody overnight, membranes were washed and incubated with secondary HRP-conjugated AffiniPure antibodies (Jackson ImmunoResearch) for 1 h at room temperature and subsequently incubated with Enhanced Chemiluminescence Detection solution (Amersham).

### Senescence Associated β-Galactosidase (SA-β-Gal) staining

SA-β-Gal staining was performed using the Senescence β-Galactosidase Staining kit (Cell Signalling), following the manufacturer instructions. Briefly, cells were fixed at RT for 15 min with a 2% formaldehyde, washed with PBS and incubated overnight at 37 °C with the staining solution containing X-gal in N,N-dimethylformamide (pH 6.0 adjusted with HCl). The next day cells were washed 3x with PBS for 2 min, and finally PBS was placed over the cells for imaging. Pictures were taken using a Wide Field Zeiss Axio Observer 7. For tissue cryosections this method was repeated, however the tissue was incubated with x-gal for 4–6 h.

### Immunohistochemistry and Immunofluorescence

For Immunohistochemistry, tumors and organs were extracted and put in 10% neutral buffered formalin (4% formaldehyde in solution). They were then transferred to 70% ethanol. The samples were then embedded in paraffin by the Histopathology Core Facility of CRUK Cambridge Institute, and sent for processing at CNIO, where they were stained using Phospho-Rb (Ser807/811), and Ki-67 (D3B5). Digital image analysis was performed using HALO and the CytoNuclear v2.0.9 imaging module (Indica Labs, Albuquerque, USA).

Immunofluorescence was performed on samples previously placed in OCT and frozen. 10 µm slices were made using a Leica CM3050 S cryostat. Briefly, samples were washed twice with PBS, fixed in 4% PFA solution for 10 min, washed again with PBS, and then permeabilized with 0.25% Triton X-100. Samples were then blocked for 1 h using 2% normal donkey serum. In the same blocking solution, a 1/500 dilution of the primary antibody for Ki-67 Rabbit host (ICH-00375) from Bethyl Laboratories. After incubation overnight at 4 °C, slides were washed twice in PBS and anti-rabbit secondary was incubated for 2 h at RT. Alexa Fluor® 488 AffiniPure Donkey Anti-Rabbit IgG was used (711-545-152) from Jackson Immuno Research. Slides were then mounted using fluoromount-G (0100-01) from SouthernBiotech and imaged using the Leica SP5 confocal microscope using a 20× HCX PL APO 0.5 NA dry objective or a 40× HCX PL APO 1.3 NA oil immersion objective. Images were analyzed with LAS AF Lite (Leica).

### Statistical Analysis

All analysis was performed unblinded. Statistical analyses were performed as described in the figure legend for each experiment. Statistical significance was determined by Student’s t tests (two-tailed) using Prism 9 software (GraphPad) as indicated. A p-value below .05 was considered significant and indicated with asterisk: *p < .05, **p < .01, ***p < .001, and ****p < .00001.

## Acknowledgements

We thank the members of the Fruk, Munoz-Espin, and Bohndiek groups for constant support in and out of the lab. We would also like to thank Gemma Cronshaw and all of her team in the biological resource unit (BRU) of the Cancer Research UK Cambridge Institute for their support with the animal models and day to day maintenance. Further we would like to thank Andrew Mason, Duncan Howe, and Pete Gierth of the Yusuf Hamied Department of Chemistry NMR facility. As well as, Asha Boodhun, Roberto Canales, and Dijana Matak-Vinkovic of the Department of Chemistry Mass Spectrometry.

## Author contributions

Conceptualization: AGB, DME, LF Methodology: AGB, DME, LF ELB, JJ

Investigation: AGB, HO, MH, EG, DM, DME, CS, ELB, MG, ABP, TRE, AV, HFG

Image Analysis: AGB, JJ Supervision: DME, LF, SEB Writing: AGB, DME, LF

Review & Editing: AGB, ABP, DME, LF, SEB, ELB, HO, MH, EG, DM, CS, MG, TRE, AV, HFG

## Competing interest: Data and materials availability

AGB, DME and LF have filed a patent for the present technology. All data associated with this study are present in the paper, Materials and Methods or Supplementary Materials.

## Extended Data for

**Extended Data Figure 1.**
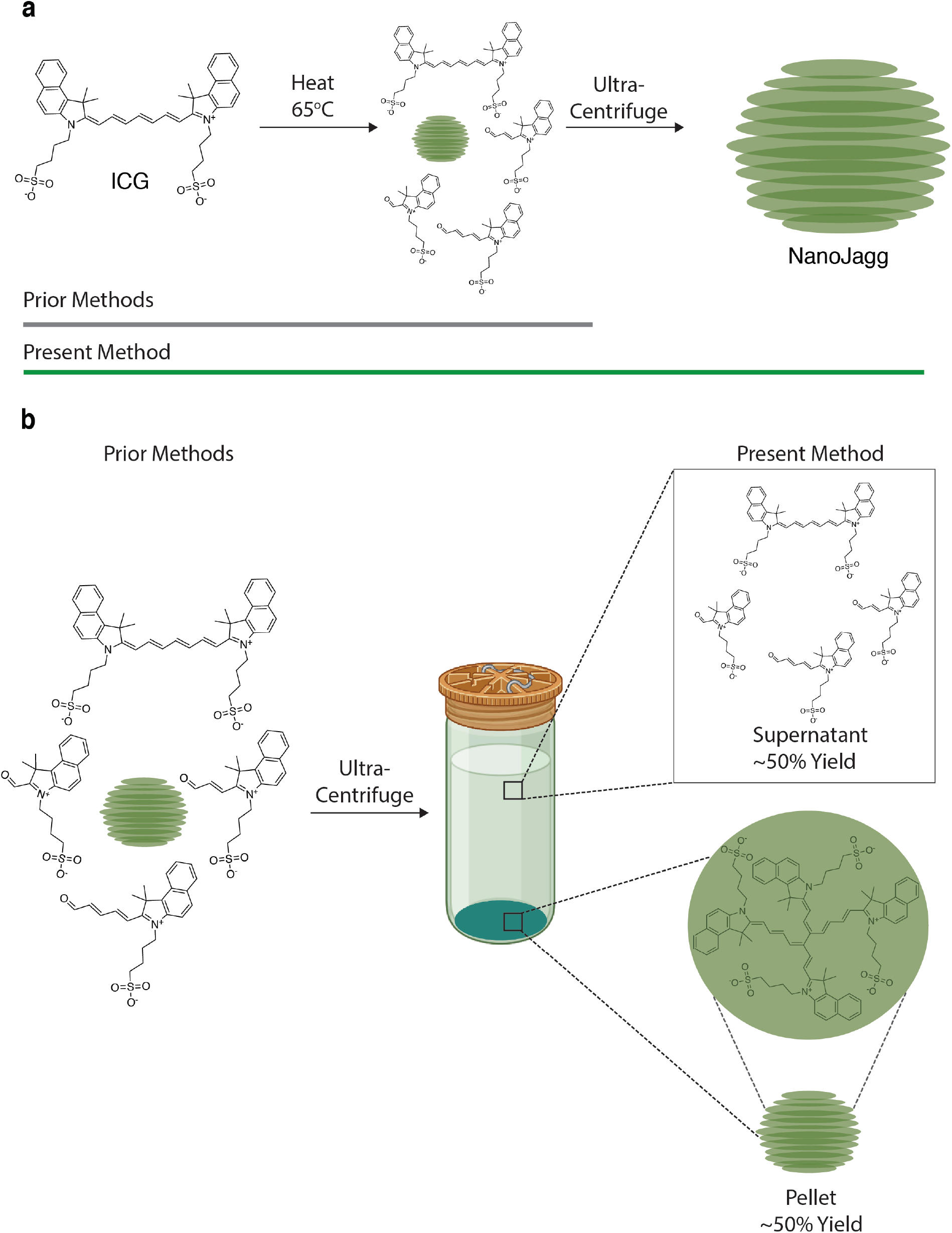
Preparation of NanoJaggs and some of the side products obtained in previously reported methods. **(a)** Prior reported methods do not report purification step. In addition, solutions were not fully characterized. Introduction of centrifugation step results in pure NanoJaggs. (**b**) Some of the side products identified in the reaction mixture. After centrifugation, a dark green pellet obtained represents ∼50% by mass yield of the reaction and contains NanoJaggs, J-aggregate nanoparticles, composed of a dimer of ICG as opposed to what was previously believed to be a monomer.

**Extended Data Figure 2.**
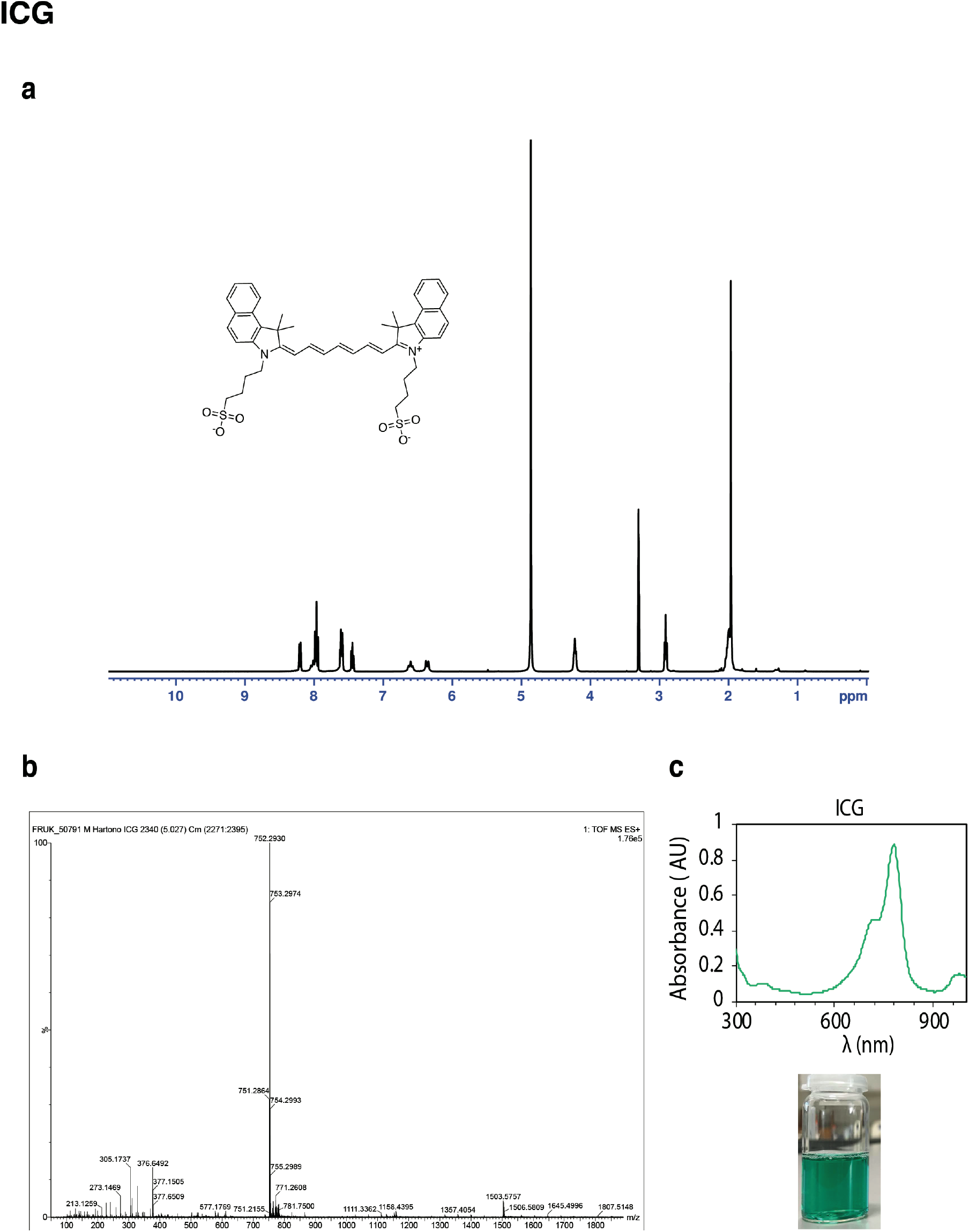
Indocyanine Green Dye (ICG) characterization. (**a**) ^1^H NMR spectrum, (**b**) LC-MS/MS, and (**c**) UV-Vis of green ICG solution. To obtain the NMR spectrum, 10 mg of ICG is dissolved in deuterated methanol (CD3OD), while mass spectrometry was performed using ICG in methanol.

**Extended Data Figure 3.**
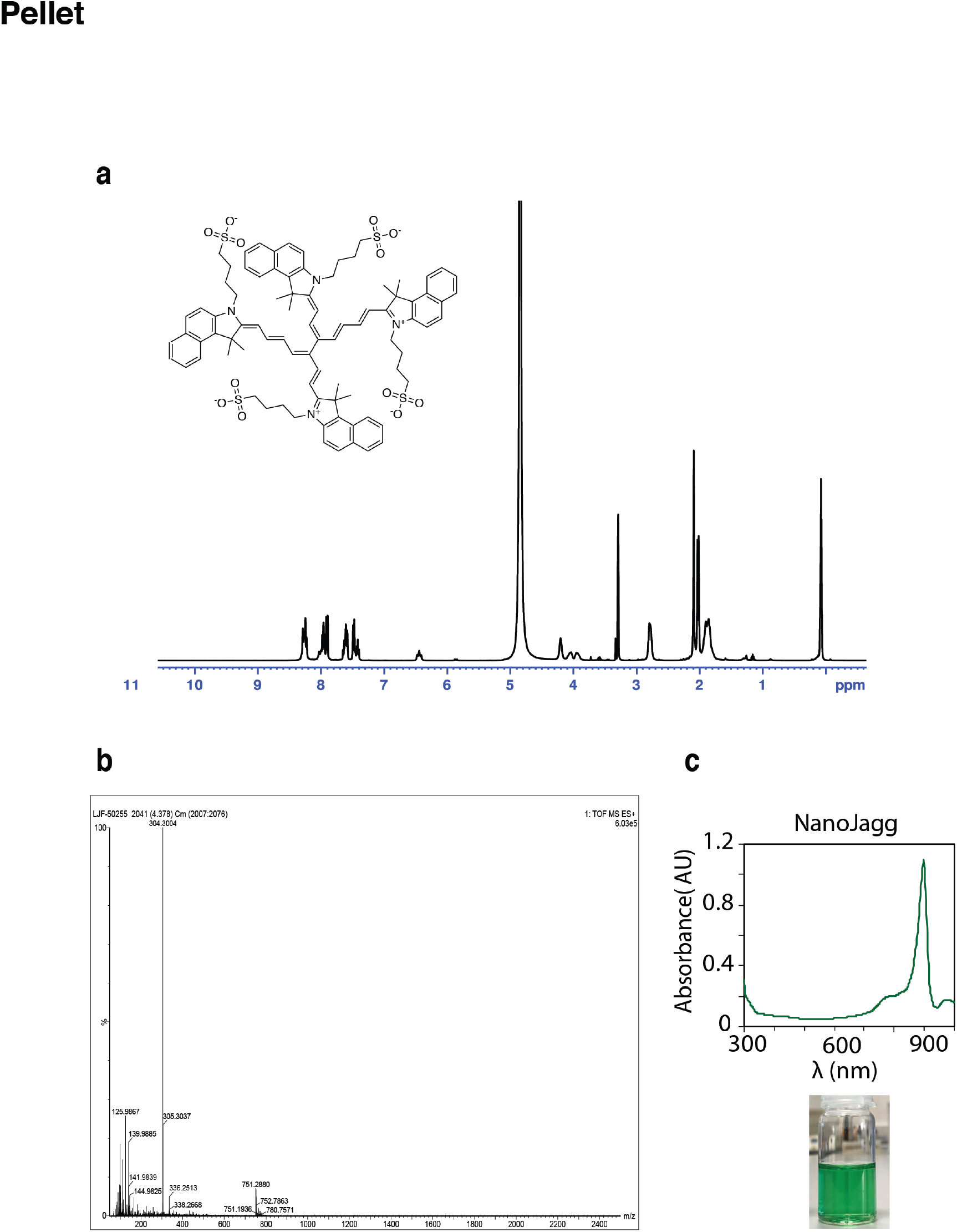
Characterization of NanoJagg pellet. (**a**) ^1^H NMR spectrum, (**b**) LC-MS/MS data, and (**c**) UV-VIS spectrum. ^1^H NMR was obtained using 10 mg of the pellet dissolved in deuterated methanol (CD3OD) while pellet dissolved in methanol was used for mass spectrometry.

**Extended Data Figure 4.**
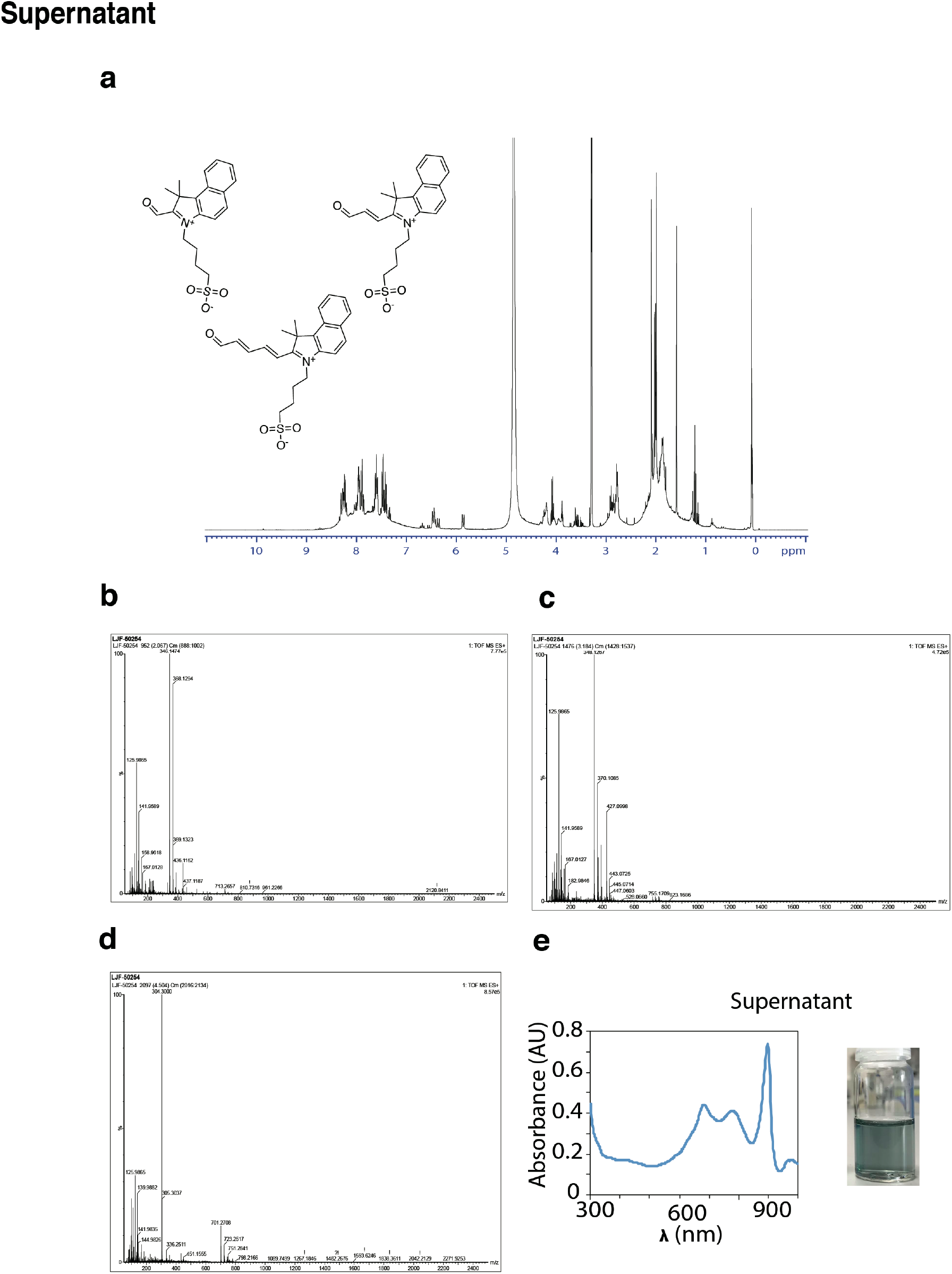
Characterization of the supernatant. (**a**) ^1^H NMR spectrum, (**b**, **c**, **d**) LC-MS/MS data, and (**e**) UV-VIS spectrum including the photo of the supernatant. ^1^H NMR was obtained using deuterated methanol (CD3OD).

**Extended Data Figure 5.**
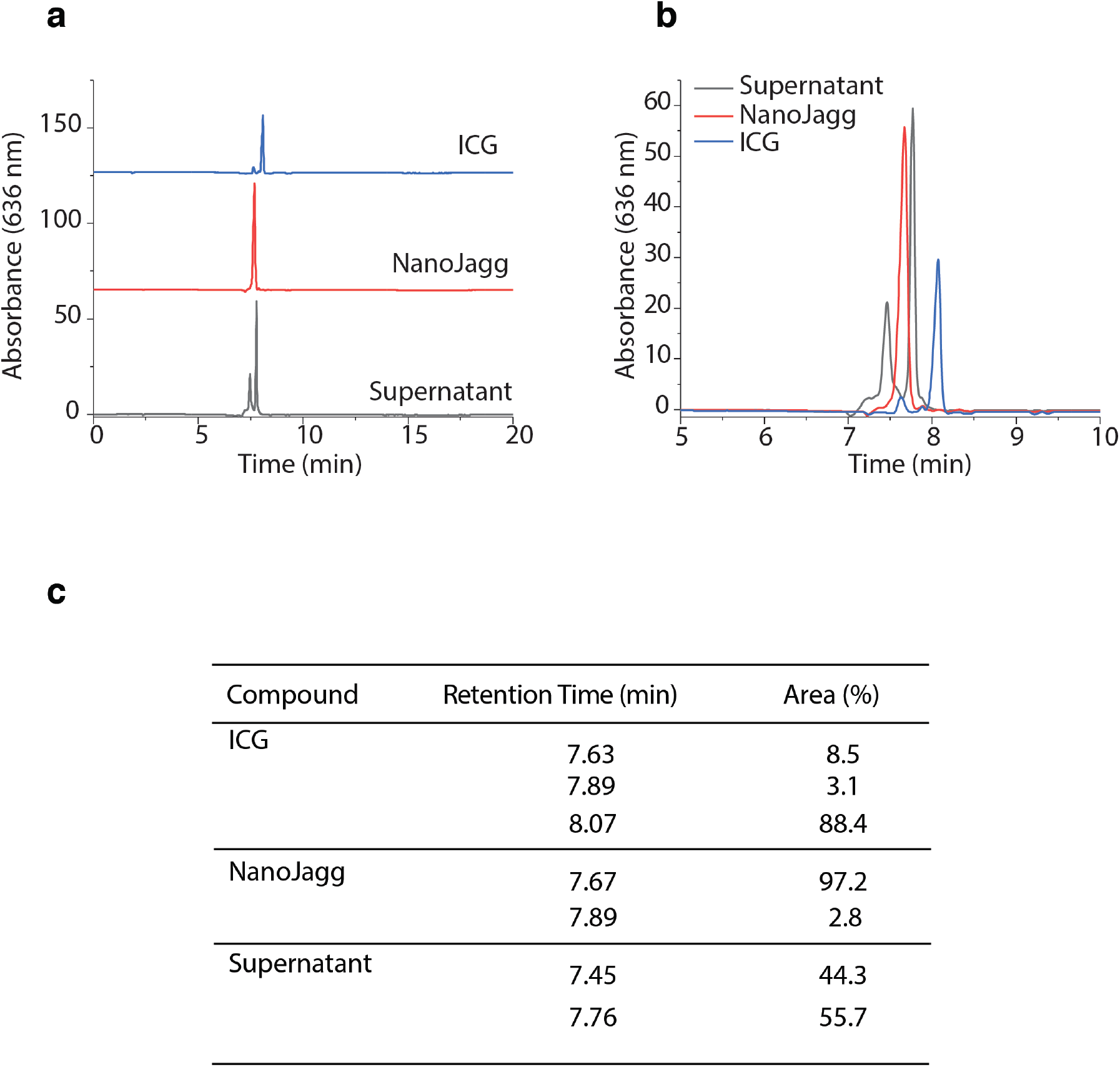
HPLC traces and retention times of ICG, NanoJaggs and supernatant. Individual (**a**) and combined (**b**) HPLC traces of ICG, NanoJaggs and supernatant illustrating the differences in retention times. (**c**) Retention times of individual samples and percentage area obtained calculating the peak area of each individual trace.

**Extended Data Figure 6.**
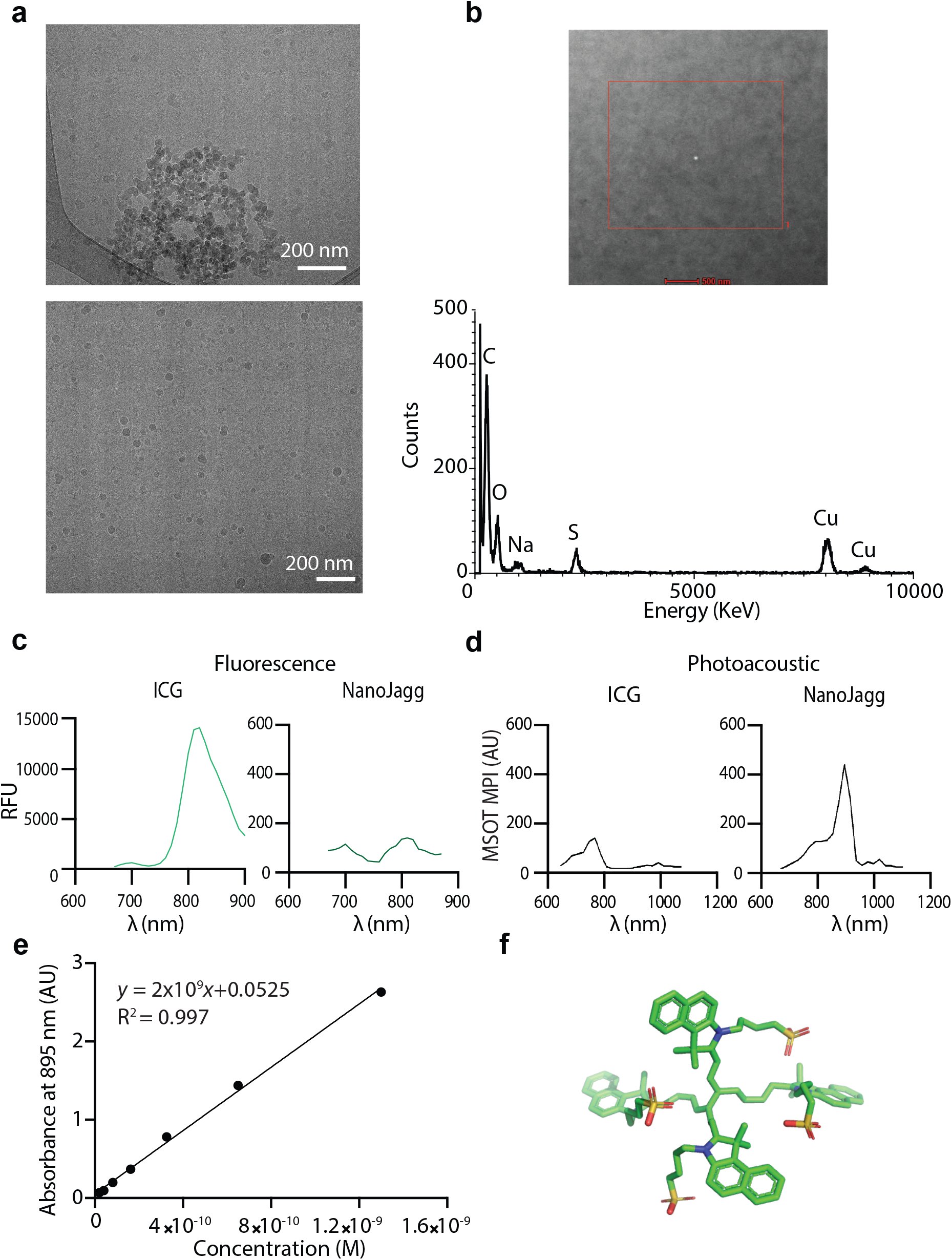
Chemical stability in organic solvents, and photoacoustic properties of ICG and NanoJaggs. (**a**) Cryo TEM images of NanoJaggs. (**b**) Energy dispersive X-ray analysis (EDX) demonstrates presence of (C) carbon, (O) oxygen, (Na) sodium, (S) sulfur, and (Cu). (**c**) Fluorescence emission spectrum excitation at λ_ex_ = 633 nm and (**d**) photoacoustic spectrum of ICG and NanoJaggs. Both solutions were normalized to a peak absorbance of 1AU prior to analysis. (**e**) Absorbance vs concentration calibration curve to estimate the extinction coefficient of NanoJagg at the peak absorbance of 895 nm. X-axis (Concentration) is referring to calculated amount of NanoJagg nanoparticles. (**f**) Structure of the dimer of indocyanine green created in PyMOL.

**Extended Data Figure 7.**
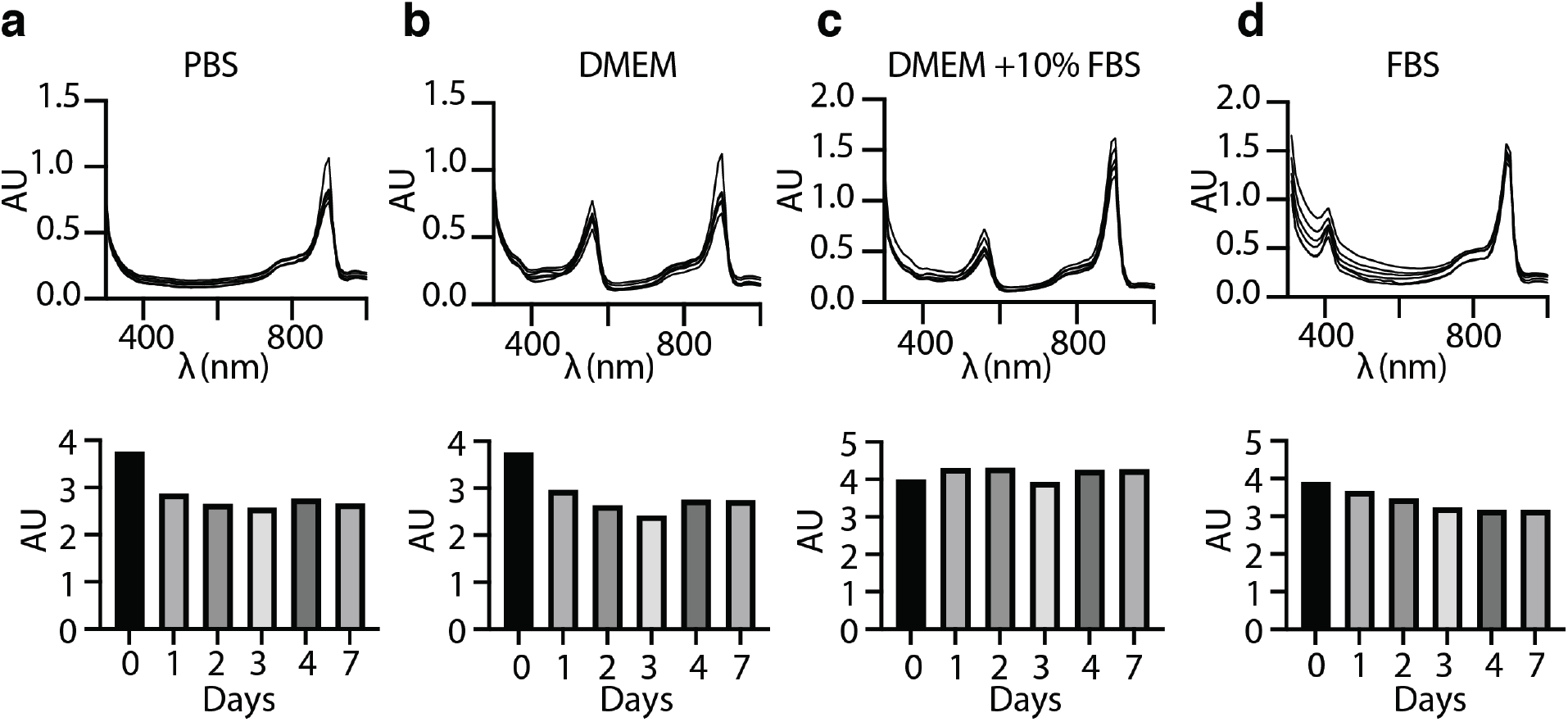
Physiochemical stability of NanoJaggs in biological media. (**a**) Phosphate buffer pH 7.5, (**b**) Dulbecco’s Modified Eagle Medium (DMEM), (**c**) DMEM + 10% Fetal Bovine Serum (FBS) and (**d**) 100% FBS. NanoJaggs were incubated over 1 week at 37 °C and data was obtained by monitoring absorbance spectra at 895nm and 780nm.

**Extended Data Figure 8.**
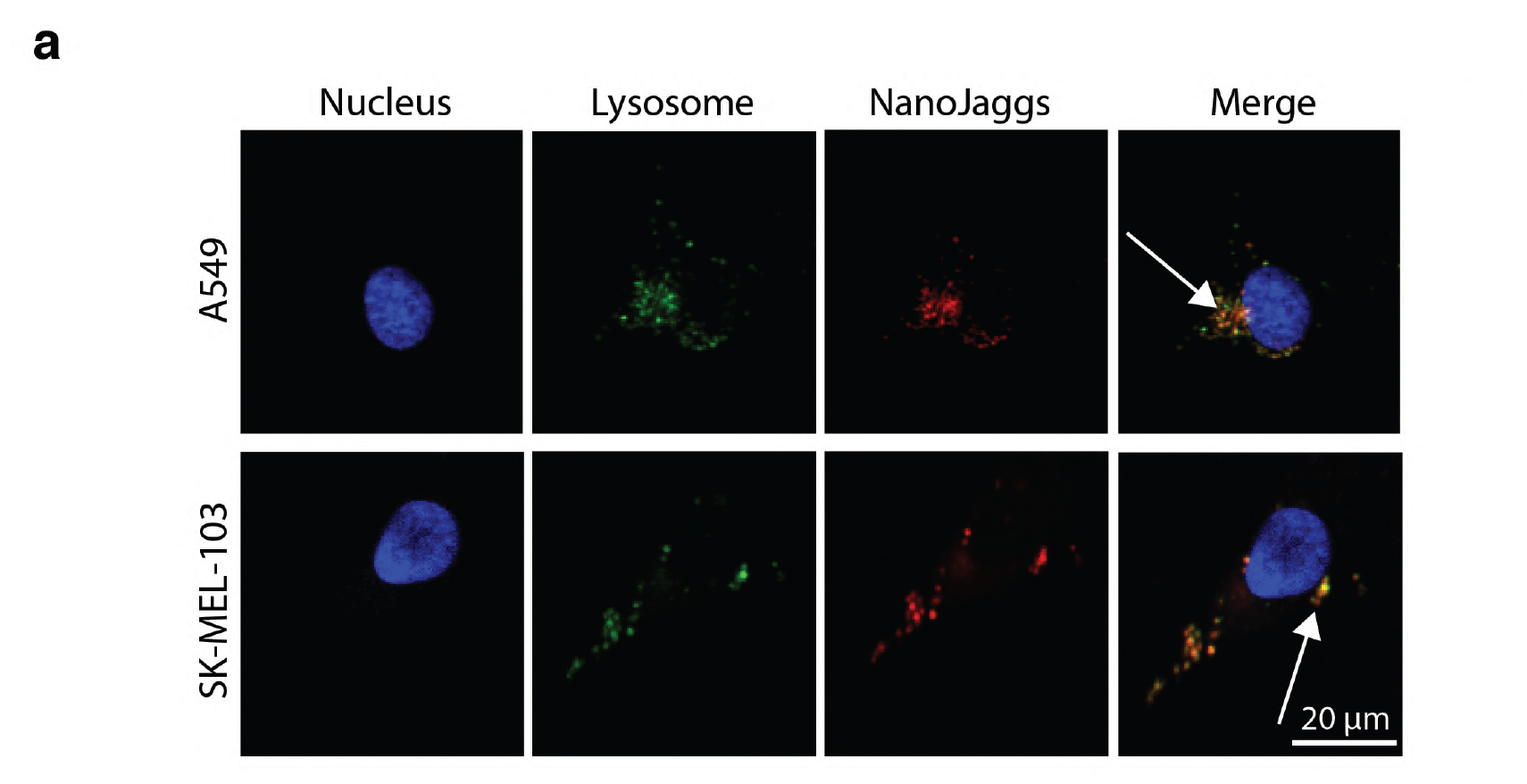
NanoJagg accumulation in the lysosomes of cancer cells. (**a**) NanoJaggs were added at 10 µg/mL to control lung cancer (A549) and melanoma cancer (SK-MEL-103) cells. Nucleus was stained using Hoechst stain, lysosome with LysoTracker Green, NanoJaggs excited at 633nm and emission between 680-720nm. Confocal images obtained using Zeiss Axio Observer Z1.

**Extended Data Figure 9.**
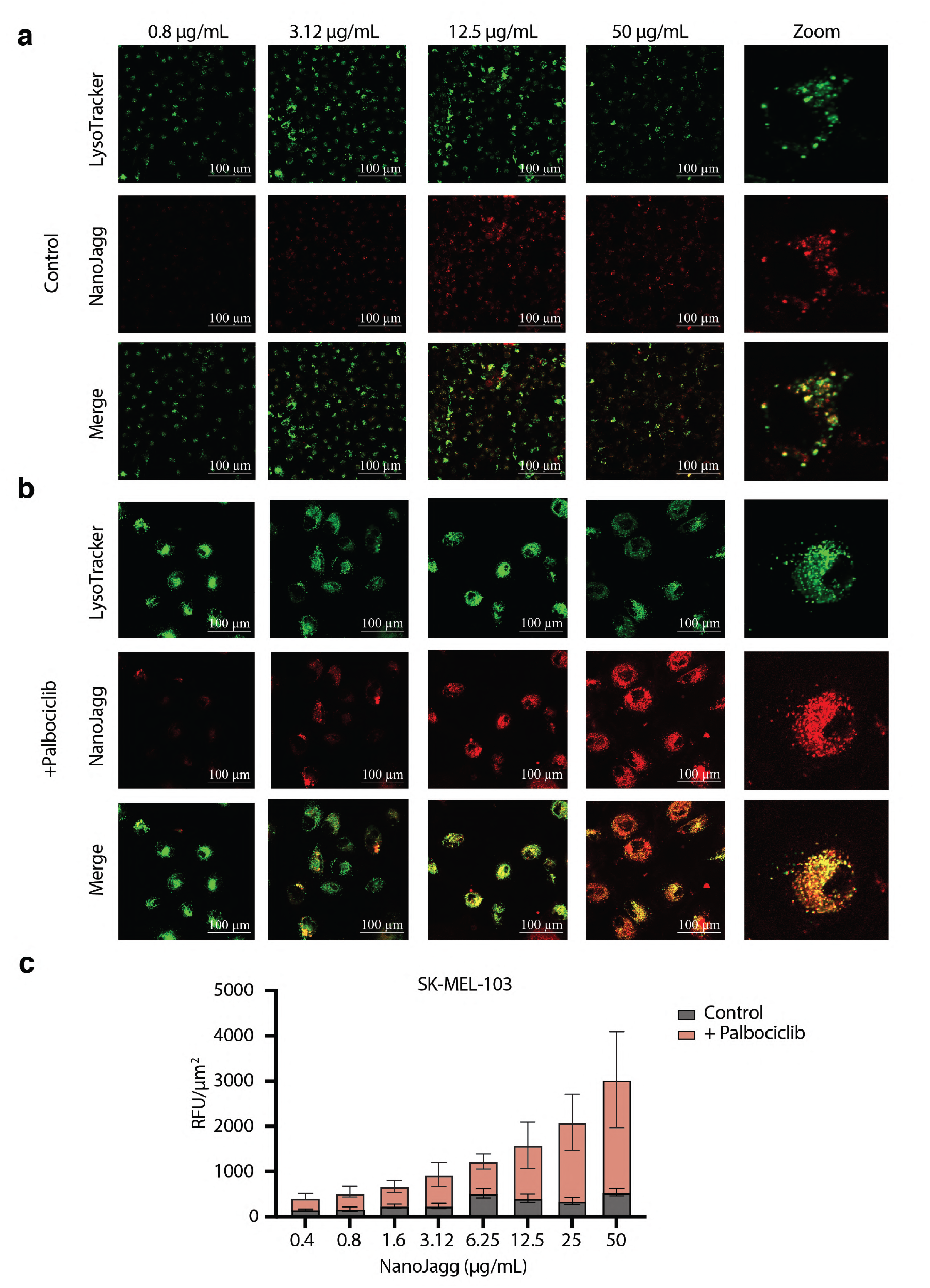
NanoJagg accumulation in the lysosomes of cancer (Control) and senescent (+ Palbociclib) cells. (**a**) NanoJaggs are taken up by SK-MEL-103 cancer cells and can be found in lysosomes. (**b**) Senescent SK-MEL-103 cells induced by 7-day treatment with 5 µM of Palbociclib show increased uptake into lysosomes. (**c**) Quantification of NanoJagg indicate significantly higher accumulation in senescent cells (p<0.001). Data represent mean ± SD, and a Two tailed t test was used to calculate the significance (*p < .05, **p < .01, ***p < .001, and ****p<0.0001). Lysosome was stained with LysoTracker Green, and NanoJaggs excited at 633nm and emission range 680-720nm. Confocal images were obtained using Zeiss Axio Observer Z1.

**Extended Data Figure 10.**
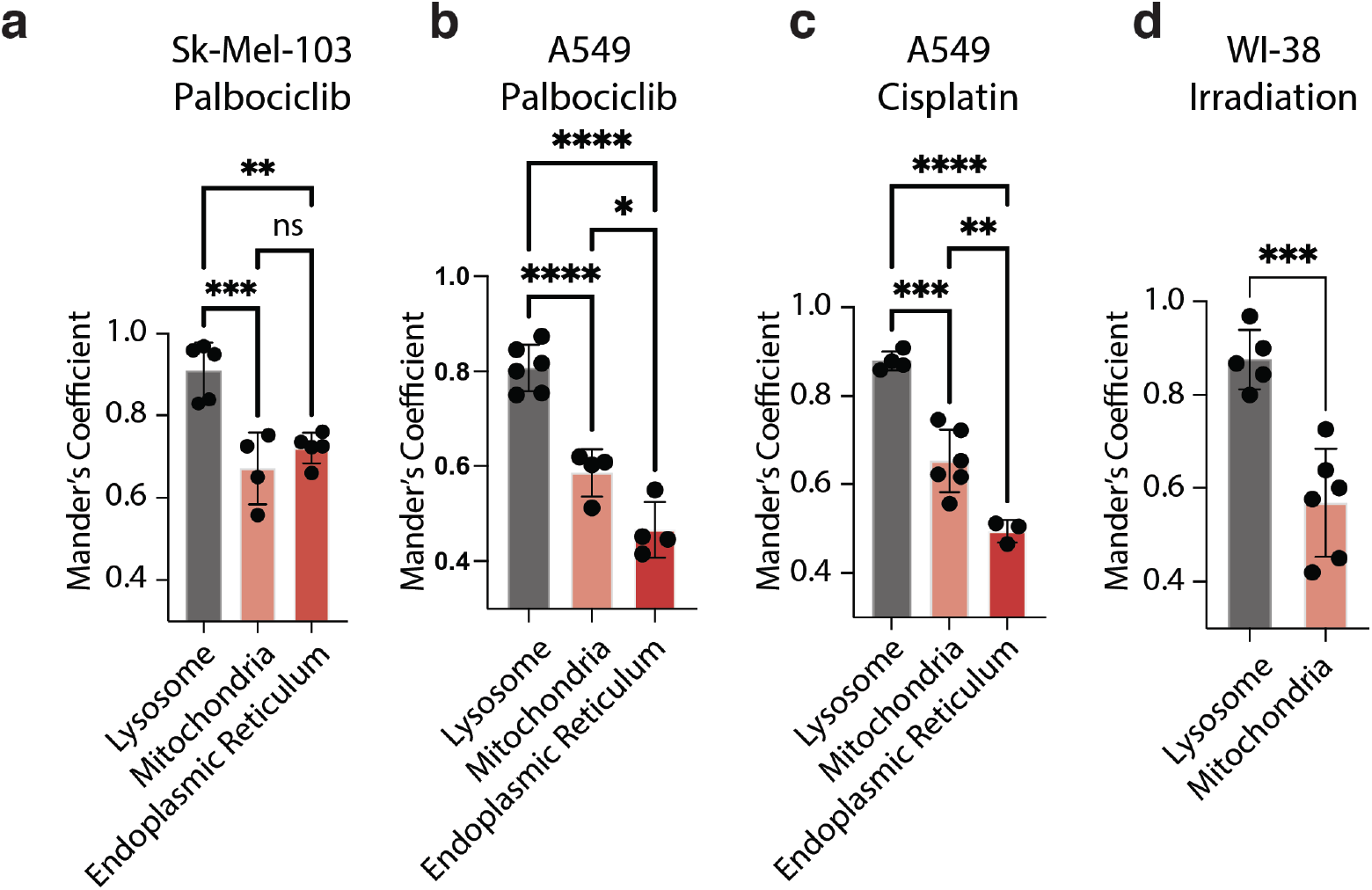
Quantification of NanoJagg colocalization in lysosomes, mitochondria and endoplasmic reticulum using Mander’s coefficient. Senescent cells were obtained using different cell treatment: (a) senescent SK-MEL-103 cells induced with treatment with 5 µM of Palbociclib (7 days), (b) senescent A549 cells induced with 10 µM of Palbociclib (10 days). (c) Senescent A549 cells induced with 15 µM of Cisplatin (10 days), and (d) senescent Wi-38 cells induced with 10 Gy of X-ray irradiation and then cultured for 10 days. Data is obtained using confocal images, and represent mean ± SD, and a Two tailed t test was used to calculate the significance (*p < .05, **p < .01, ***p < .001, and ****p<0.0001).

**Extended Data Figure 11.**
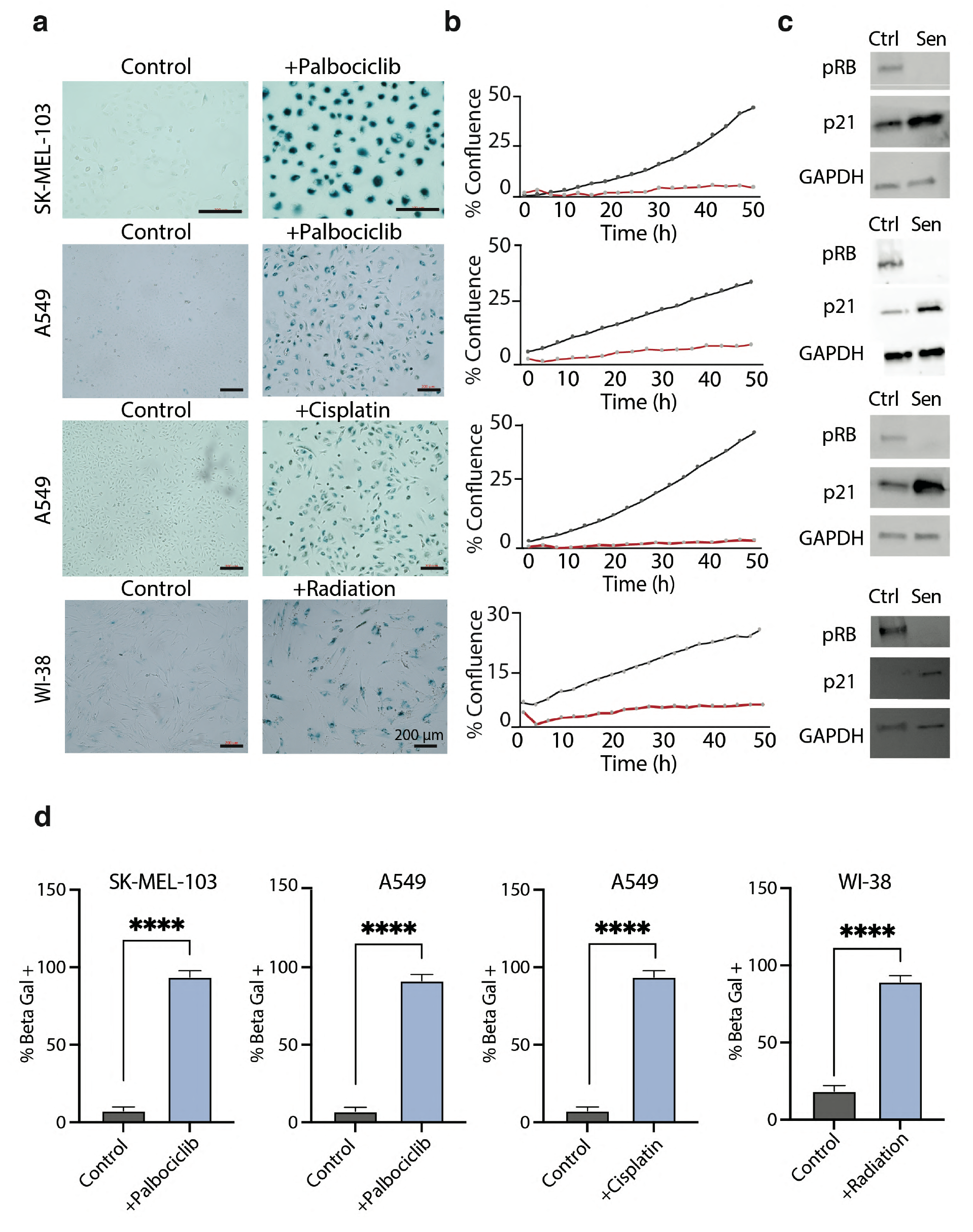
Characterization of senescent cells. Senescent cell is defined as a cell with an increased expression of β-galactosidase, reduce growth capacity, increased p21 expression and reduced expression of pRB. (**a** and **d**) Senescence-associated β-galactosidase staining using a commercial β-galactosidase staining kit. (**b**) Growth curves of control (black) and senescent (red) cells. (**c**) Western blots of cellular senescence markers, demonstrating a reduction in pRB, and increased p21 levels in senescent cells compared to controls.

**Extended Data Figure 12.**
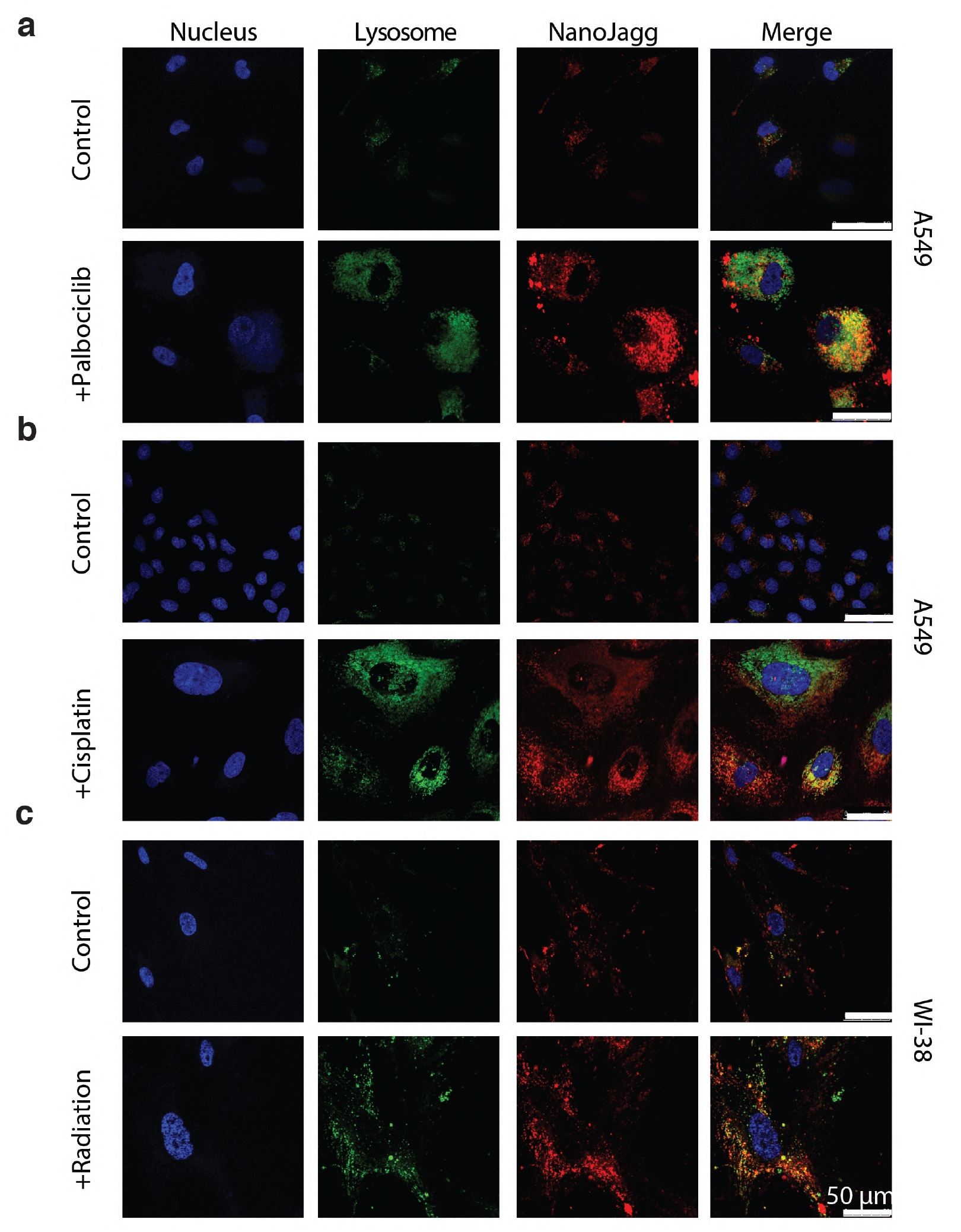
Colocalization of NanoJaggs with the lysosome of senescent cell models. (**a**) Control and senescent (+Palbociclib) A549 cells. (**b**) Control and senescent (+Cisplatin) A549 cells. (**c**) control and senescent (induced by 10 Gy X-ray irradiation) WI-38 cells. Nucleus was stained using Hoechst stain and lysosomes with LysoTracker Green. All samples were treated with 50 µg/mL NanoJaggs and were excited at 633nm and emission range 680-720nm. Confocal images were obtained on a Leica SP5 confocal microscope

**Extended Data Figure 13.**
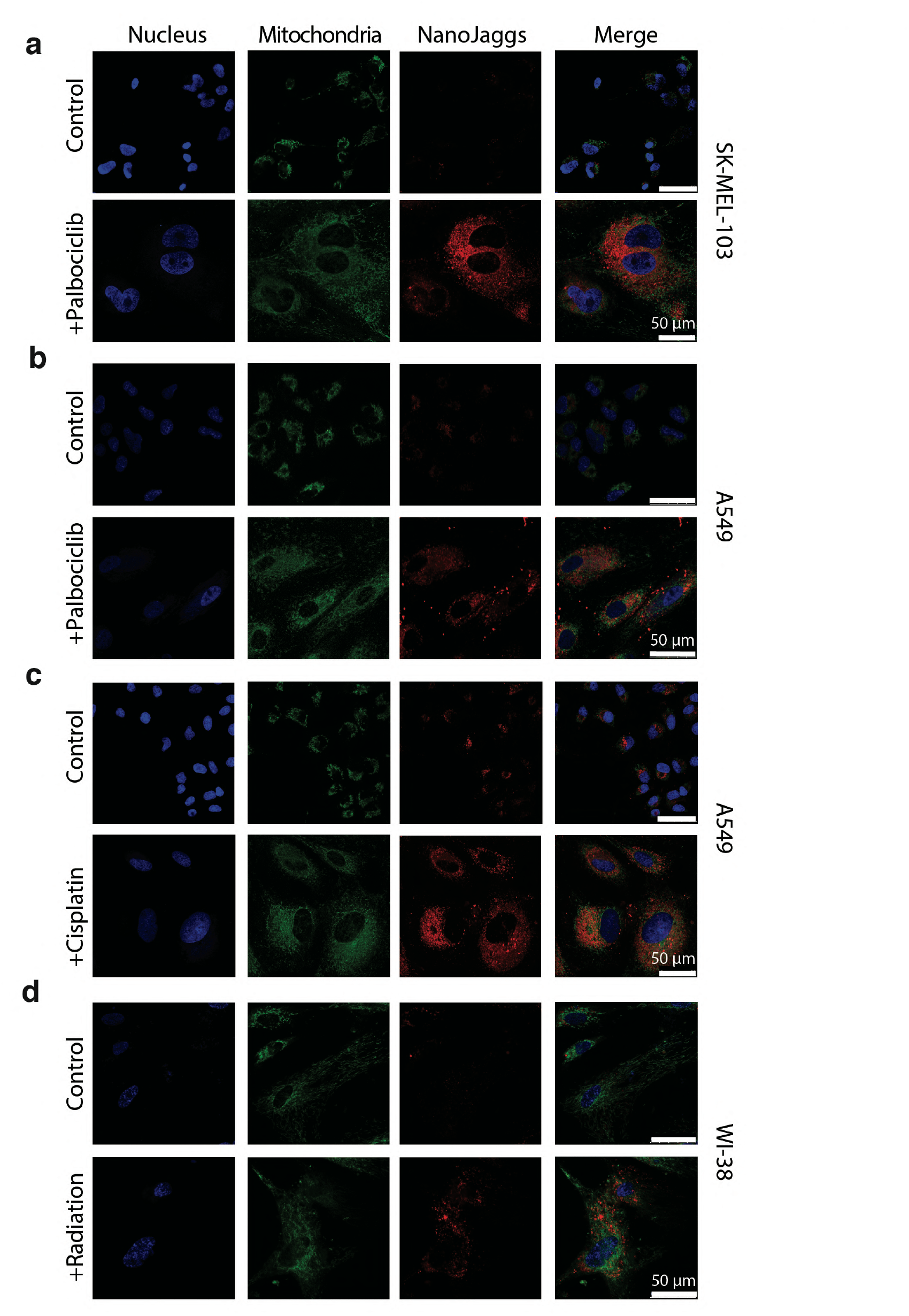
Colocalization of NanoJaggs with mitochondria. (**a**) SK-MEL-103k control and senescent cells (+Palbociclib). (**b**) A549 control and senescent cells (+Palbociclib). (**c**) A549 control and senescent cells (+Cisplatin,CDDP). (**d**) WI-38 control and senescent cells induced by X-ray irradiation. Nucleus was stained using Hoechst stain, mitochondria with MitoTracker Green. All samples were treated with 50 µg/mL NanoJaggs and were excited at 633nm and emission range 680-720nm. Confocal images were obtained on a Leica SP5 confocal microscope

**Extended Data Figure 14.**
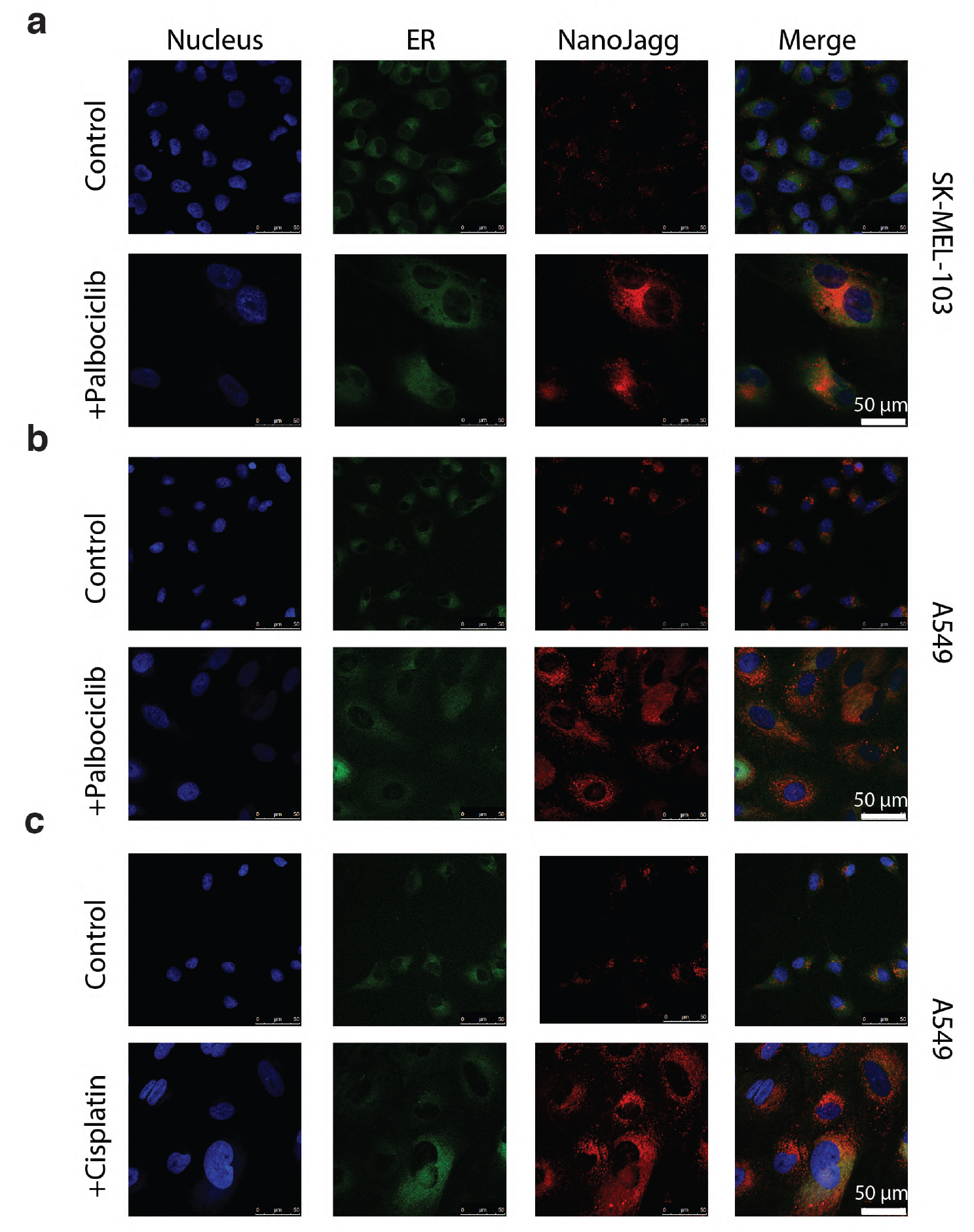
Colocalization of NanoJaggs with the endoplasmic reticulum. (**a**) SK-MELel-103 control and senescent cells (+Palbociclib). (**b**) A549 control and senescent cells (+ Palbociclib) and (**c**) A549 control and senescent cells (+cisplatin, CDDP). Nucleus was stained using Hoechst stain, and endoplasmic reticulum with ER Tracker Green. All samples were treated with 50 µg/mL NanoJaggs and were excited at 633nm and emission range 680-720nm. Confocal images were obtained on a Leica SP5 confocal microscope

**Extended Data Figure 15.**
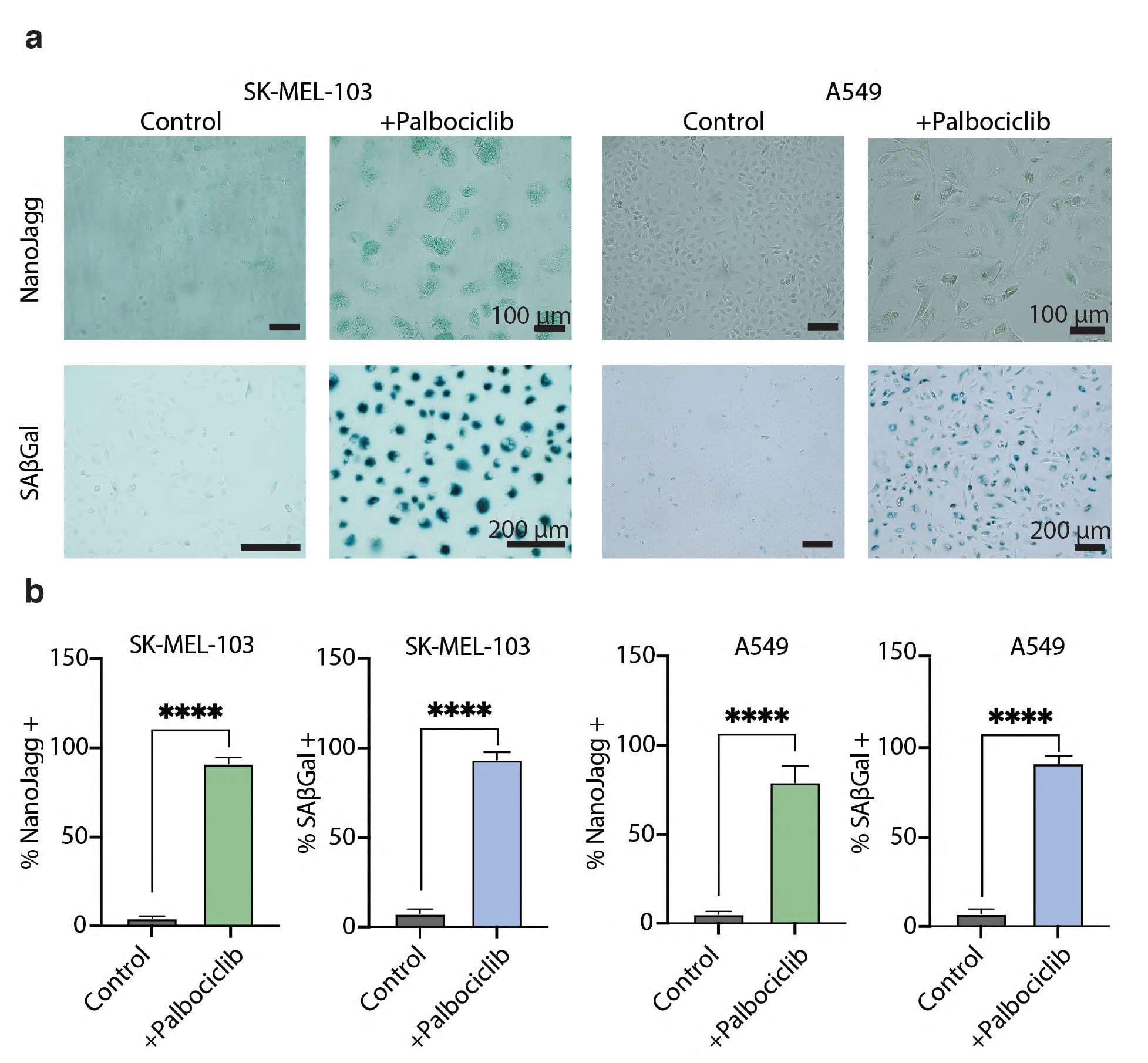
Comparison of commercial X-gal (SAβ-galactosidase) stain for senescent cells and NanoJaggs. (**a**) Control and senescent SK-MEL-103 and A549 (both +Palbociclib) cells stained with NanoJaggs and X-gal dye. (**b**) ∼90% of treated SK-MEL-103 cells were positive for NanoJaggs, and ∼95% were positive for SA-β-Gal. ∼80% of treated A459 cells were positive for NanoJaggs and ∼95% are positive to SA-β-Gal. Cells were treated with 50 µg/mL of NanoJaggs and counted to calculate % NanoJagg+ cells.

**Extended Data Figure 16.**
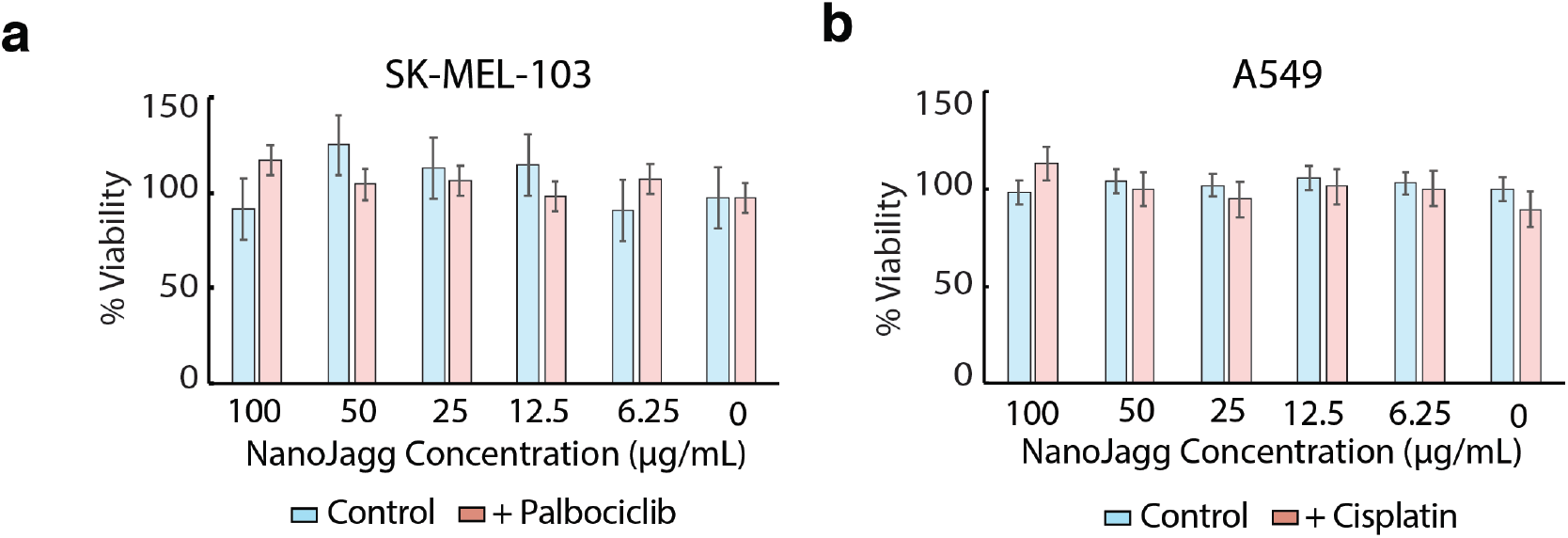
Viability of control and senescent cells treated with different concentrations of NanoJaggs. (**a**) Senescence of SK-MEL-103 was induced by 5 µM Palbociclib (7 days) and (**b**) Senescence of A549 cells was induced by 15 µM Cisplatin (10 days). Data represent mean ± SD of N=3 biological repeats. Cell viability was assessed using the CellTiter Blue assay.

**Extended Data Figure 17.**
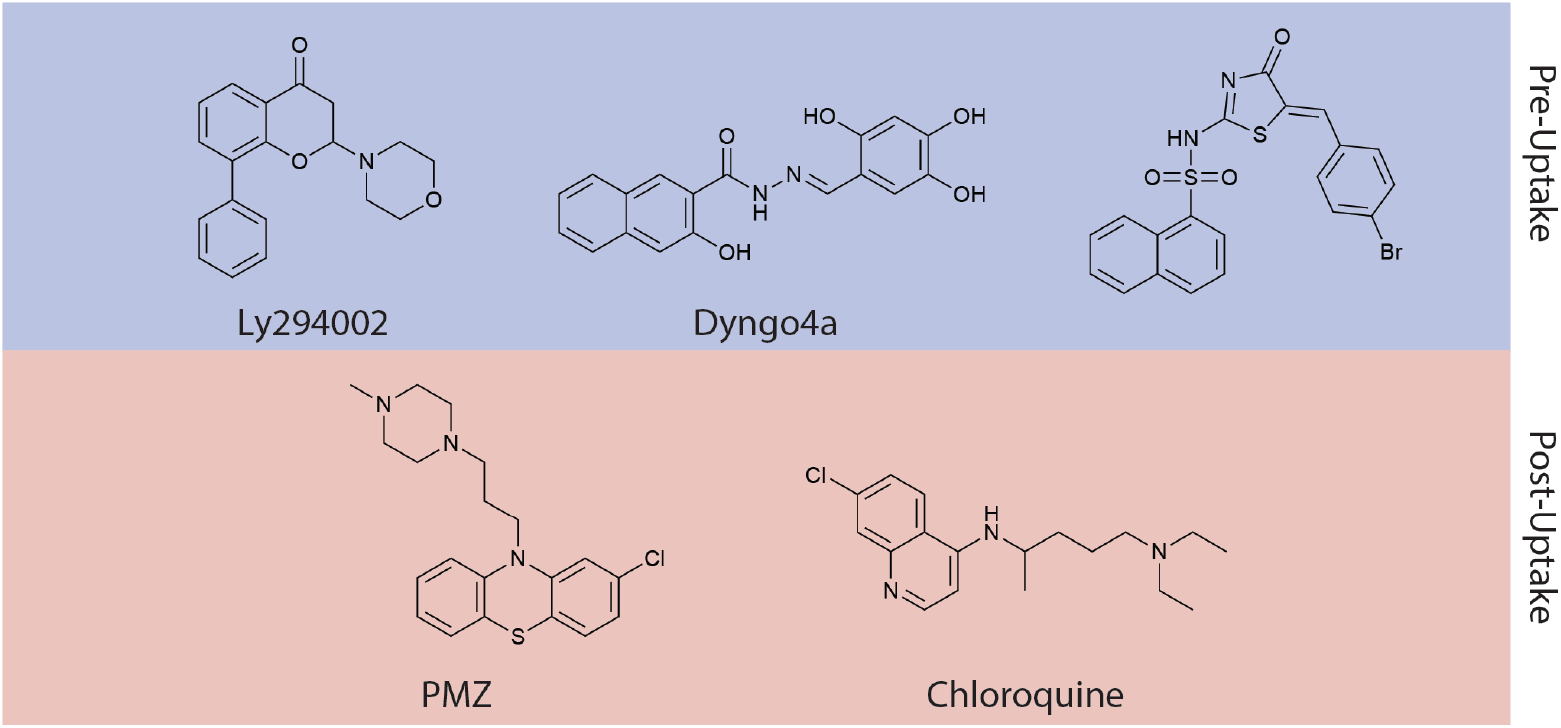
Pre-uptake and post-uptake inhibitors employed in the study. The structures of pre-uptake inhibitor Ly294002 (inhibits micropinocytosis), Dyngo4a (inhibits dynamin driven processes), and Pistop2 (inhibits the formation of clathrin coated pits.) are shown in the upper panel. Lower panel shows the structures of the post-uptake inhibitors Prochlorperazine (PMZ, inhibitor of clathrin coated pit’s fission), and chloroquine (inhibitor of autophagy).

**Extended Data Figure 18.**
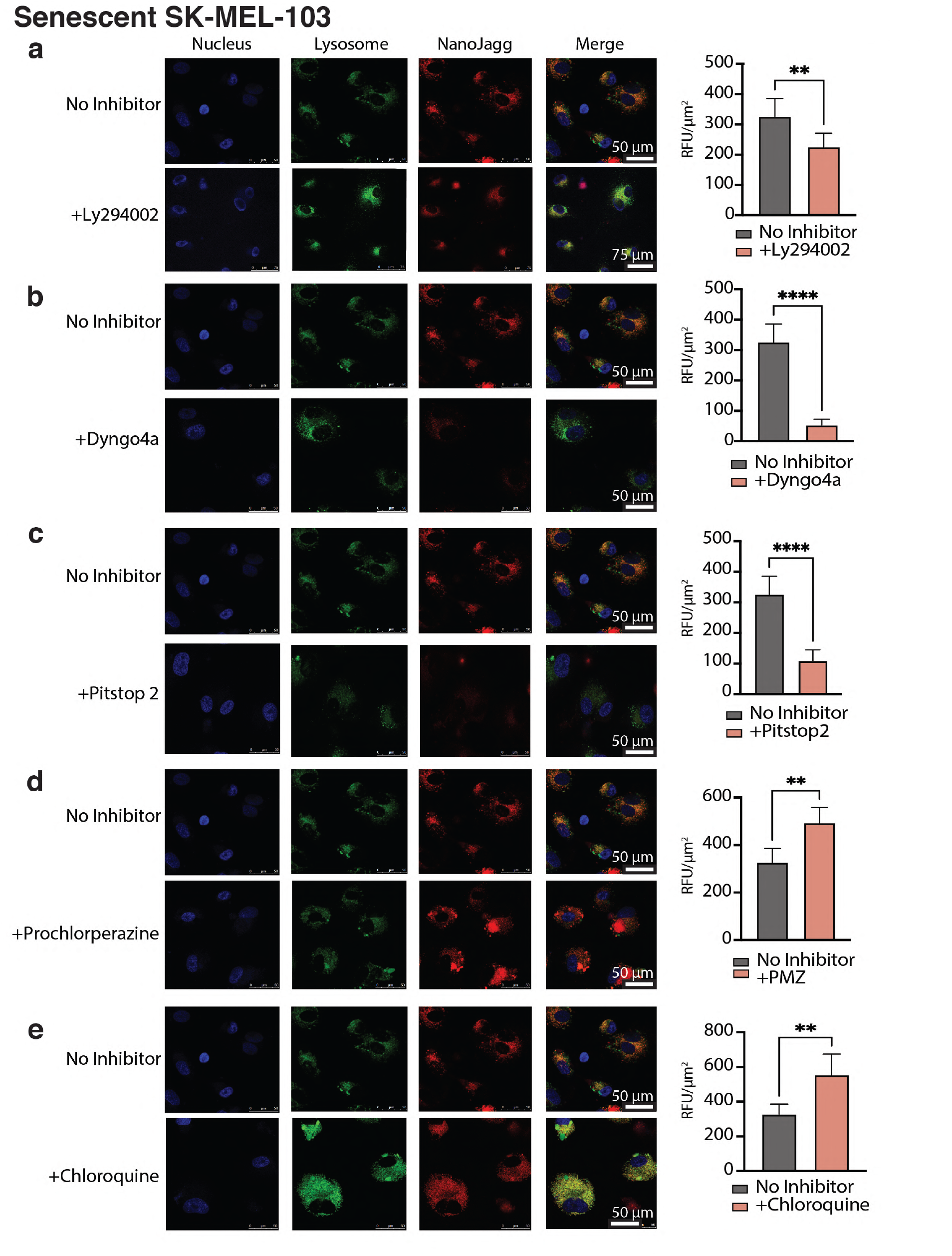
Colocalization of NanoJaggs in senescent SK-MEL-103 cells in presence of different inhibitors. Cells were treated with NanoJaggs in presence and absence of inhibitors: (**a**) LY294002 (20–50 μM, (**b**) Dyngo4a (10–30 μM), (**c**) Pitstop2 (5–15 μM), (**d**) Prochlorperazine dimaleate salt (15 μM) and (**e**) Chloroquine (50 μM). Nucleus was stained using Hoechst stain, and lysosome with LysoTracker Green. All samples were treated with 50 µg/mL NanoJaggs and were excited at 633nm and emission range 680-720nm. Confocal images were obtained on a Leica SP5 confocal microscope. Data represent mean ± SD, obtained from 3 biological replicates (N=3). Two tailed t test was used to calculate the significance (*p < .05, **p < .01, ***p < .001, and ****p<0.0001). This figure complements Figure 3 in the manuscript and provides additional data on the individual analysis for each inhibitor compared to the control done in absence of the inhibitors.

**Extended Data Figure 19.**
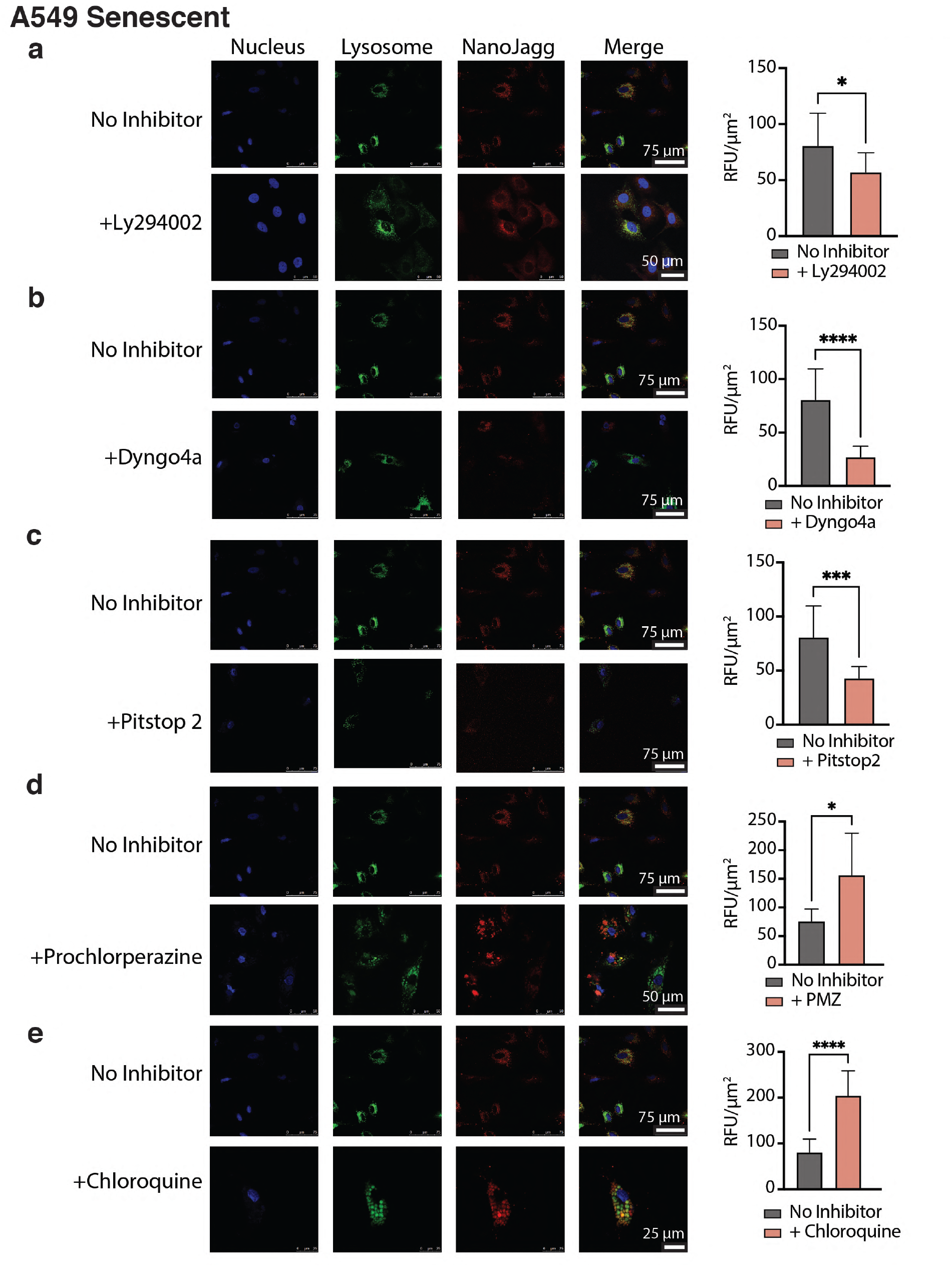
Colocalization of NanoJaggs in senescent A459 cells in presence of different inhibitors. Cells were treated with NanoJaggs in presence and absence of inhibitors: (**a**) LY294002 (20–50 μM, (**b**) Dyngo4a (10–30 μM), (**c**) Pitstop2 (5–15 μM), (**d**) Prochlorperazine dimaleate salt (15 μM) and (**e**) Chloroquine (50 μM). Nucleus was stained using Hoechst stain, and lysosome with LysoTracker Green. All samples were treated with 50 µg/mL NanoJaggs and were excited at 633nm and emission range 680-720nm. Confocal images were obtained on a Leica SP5 confocal microscope. Data represent mean ± SD, obtained from 3 biological replicates (N=3). Two tailed t test was used to calculate the significance (*p < .05, **p < .01, ***p < .001, and ****p<0.0001).

**Extended Data Figure 20.**
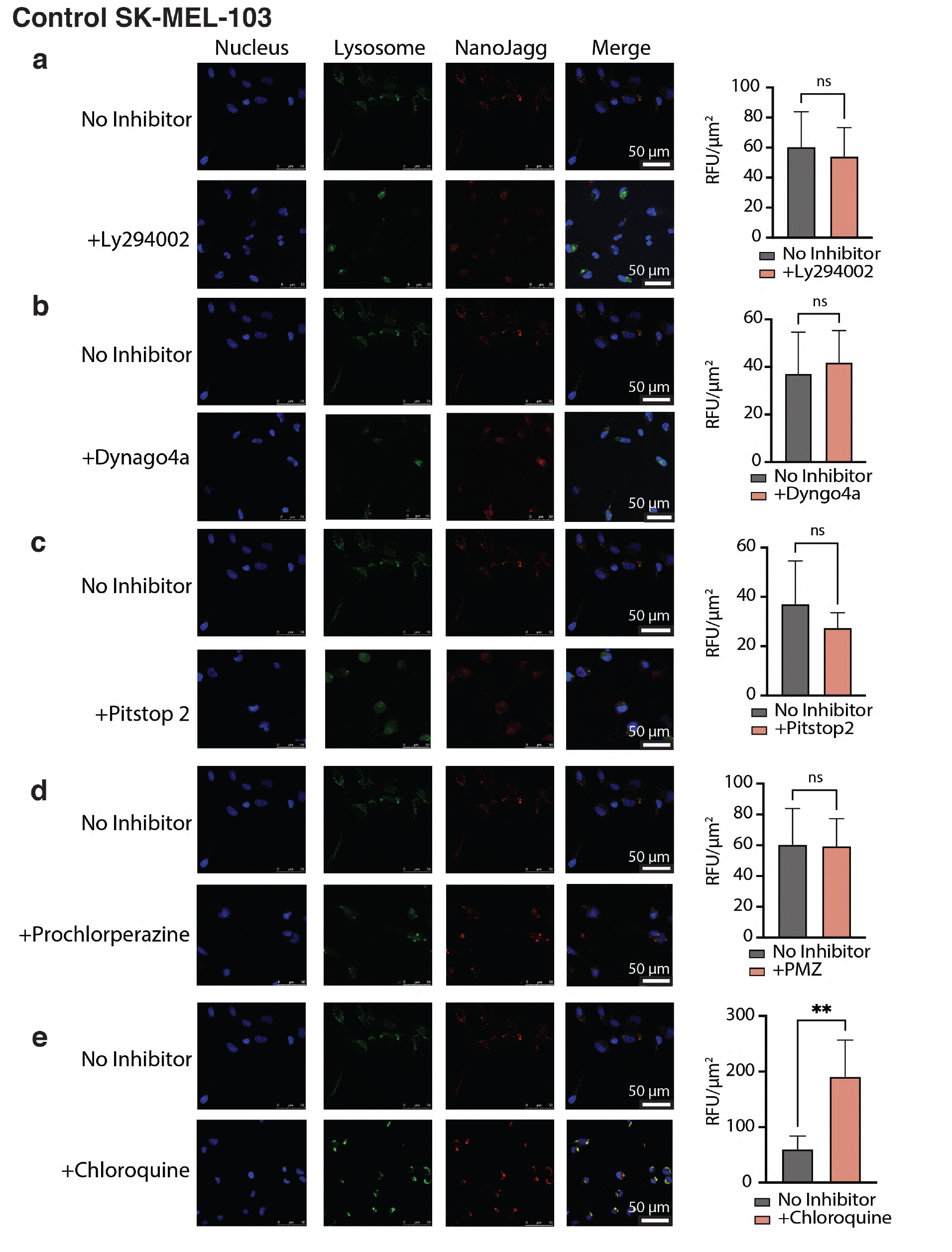
Colocalization of NanoJaggs in non-senescent SK-MEL-103 control cells in presence of inhibitors. Cells were treated with NanoJaggs in presence and absence of inhibitors: (**a**) LY294002 (20–50 μM, (**b**) Dyngo4a (10–30 μM), (**c**) Pitstop2 (5–15 μM), (**d**) Prochlorperazine dimaleate salt (15 μM) and (**e**) Chloroquine (50 μM Nucleus was stained using Hoechst stain, and lysosome with LysoTracker Green. All samples were treated with 50 µg/mL NanoJaggs and were excited at 633nm and emission range 680-720nm. Confocal images were obtained on a Leica SP5 confocal microscope. Data represent mean ± SD, obtained from 3 biological replicates (N=3). Two tailed t test was used to calculate the significance (*p < .05, **p < .01, ***p < .001, and ****p<0.0001).

**Extended Data Figure 21.**
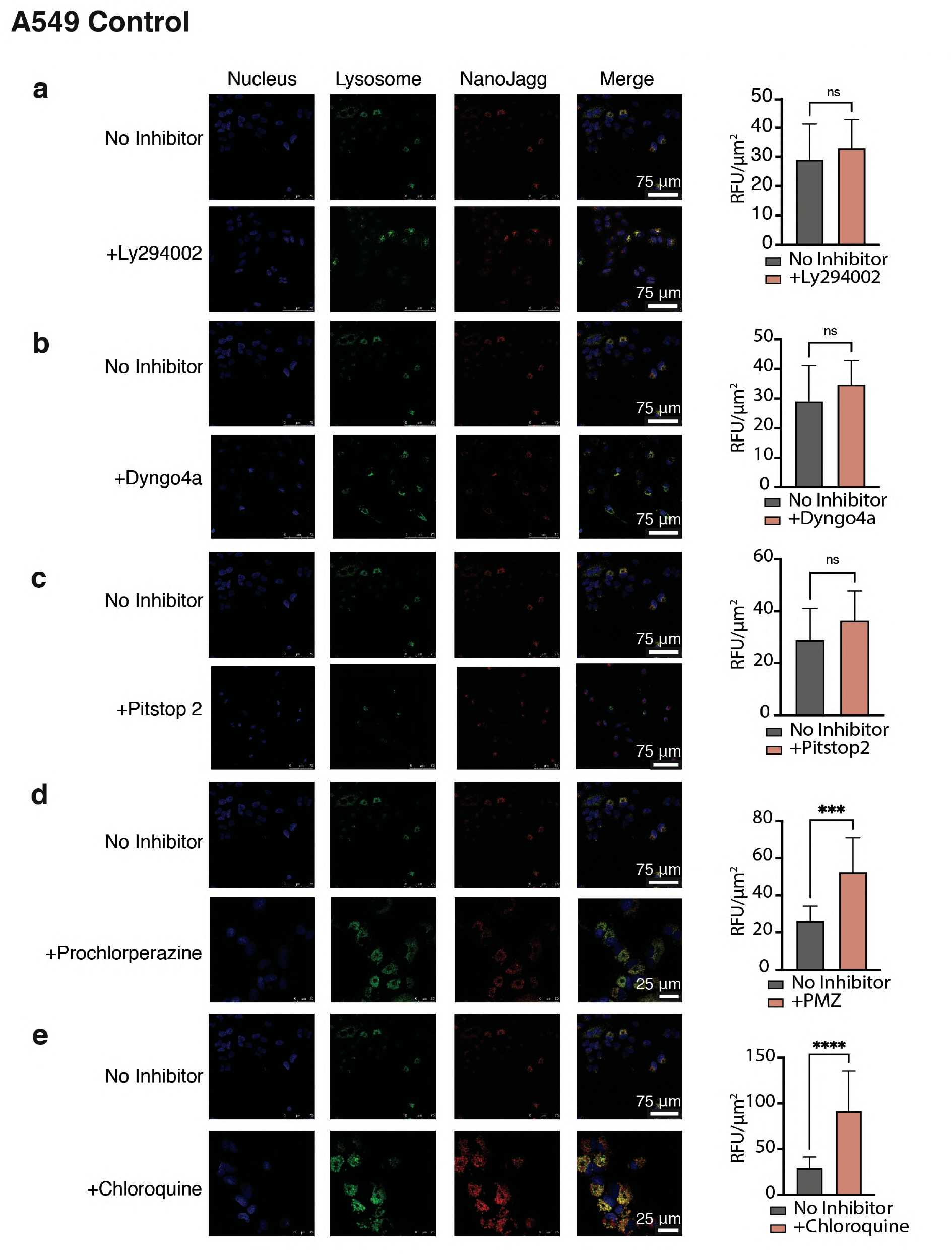
colocalization of NanoJaggs in non-senescent A459 control cells in presence of inhibitors. Cells were treated with NanoJaggs in presence and absence of inhibitors: (**a**) LY294002 (20–50 μM, (**b**) Dyngo4a (10–30 μM), (**c**) Pitstop2 (5–15 μM), (**d**) Prochlorperazine dimaleate salt (15 μM) and (**e**) Chloroquine (50 μM). (p<0.01). Nucleus was stained using Hoechst stain, and lysosome with LysoTracker Green. All samples were treated with 50 µg/mL NanoJaggs and were excited at 633nm and emission range 680-720nm. Confocal images were obtained on a Leica SP5 confocal microscope. Data represent mean ± SD, obtained from 3 biological replicates (N=3). Two tailed t test was used to calculate the significance (*p < .05, **p < .01, ***p < .001, and ****p<0.0001).

**Fig. S22.**
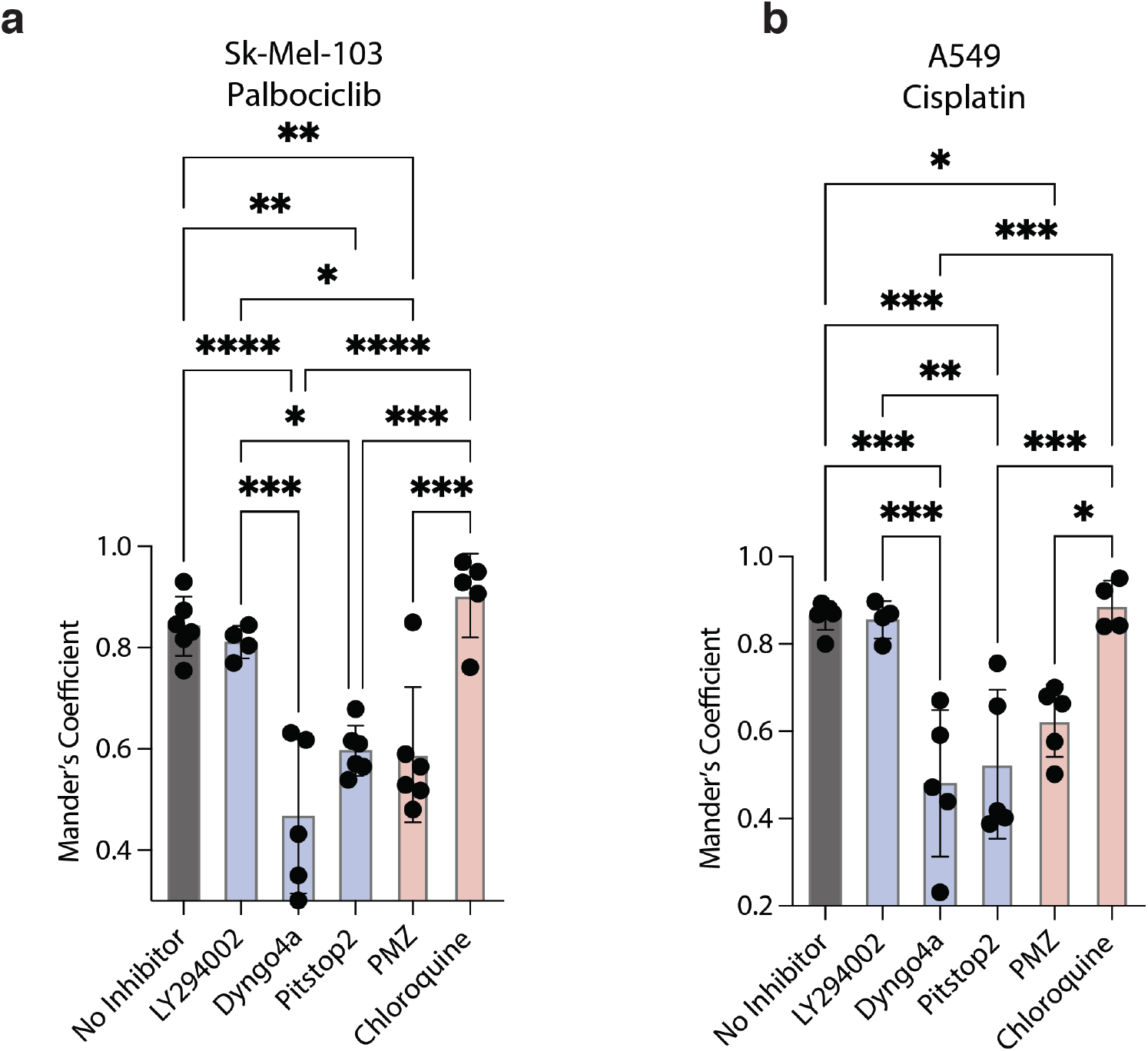
Colocalization of NanoJaggs in lysosomes of senescent cells in presence of inhibitors using Mander’s coefficient. Senescent SK-MEL-103 cells induced with 5 µM of Palbociclib for 7 days (**a**) and senescent A549 cells induced with 15 µM of Cisplatin for 10 days (**b**). Data was obtained from confocal images from Fig. S18 and S19 comparing the overlap of the lysosome and NanoJagg channel. Data represent mean ± SD, and a Two tailed t test was used to calculate the significance, the Bonferroni correction was applied (*p < .05, **p < .01, ***p < .001, and ****p<0.0001).

**Extended Data Figure 23.**
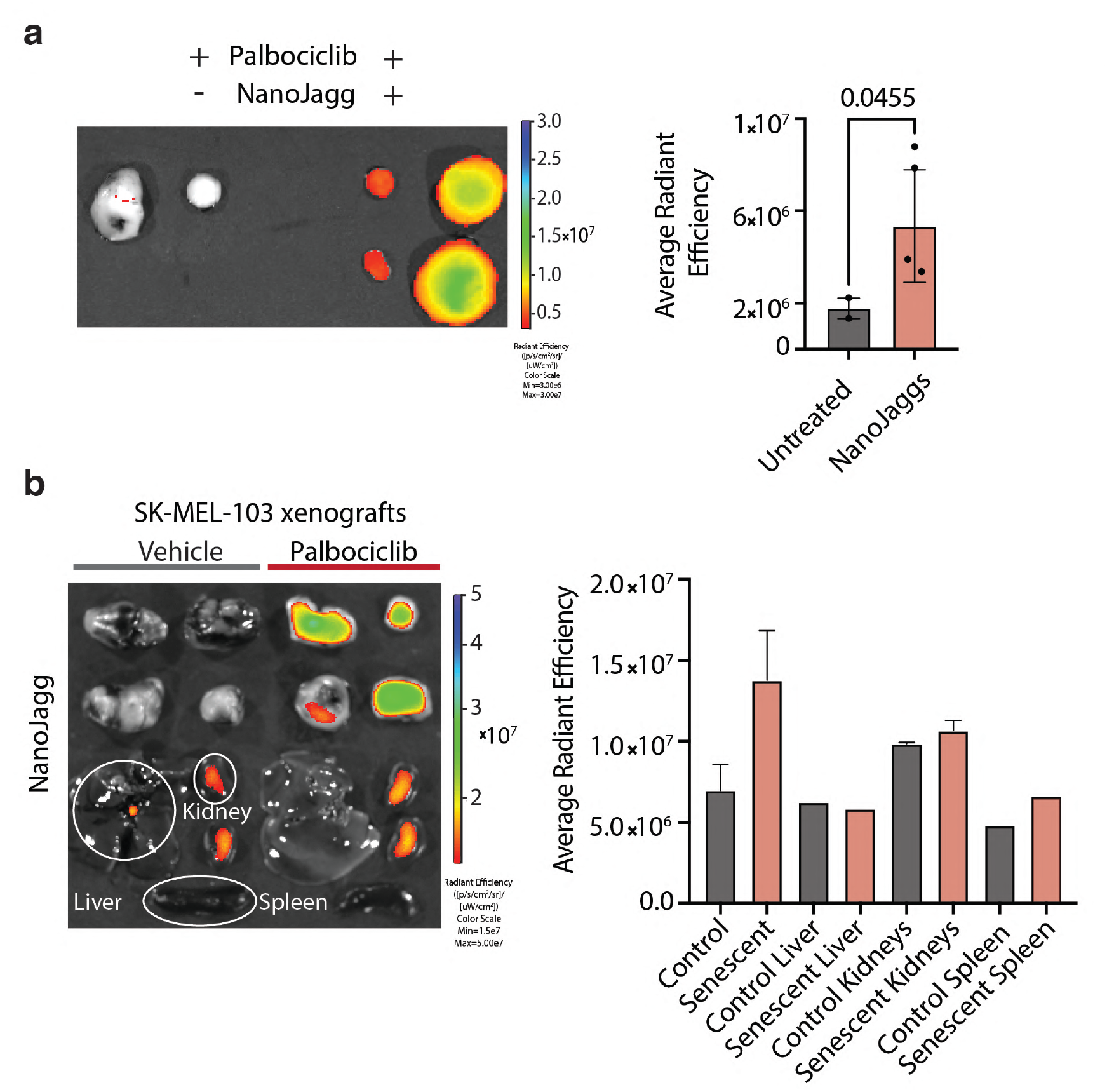
Post-in vivo assessment of NanoJaggs distribution in different organs. (**a**) Image of tumors and one set of organs from both untreated (non-senescent tumor) and Palbociclib-treated (senescent tumor) mice. (**b**) NanoJagg signal in untreated and Palbociclib treated tumors and in organs. Excitation at 610nm and emission at 700-720nm was used to monitor NanoJagg signal using an IVIS Spectrum Imaging System. Significant NanoJagg presence was detected in kidneys of both control and senescent mice indicating renal clearance. Data in graphs represent mean of the average radiant efficiency ± SD quantified from images.

**Extended Data Figure 24.**
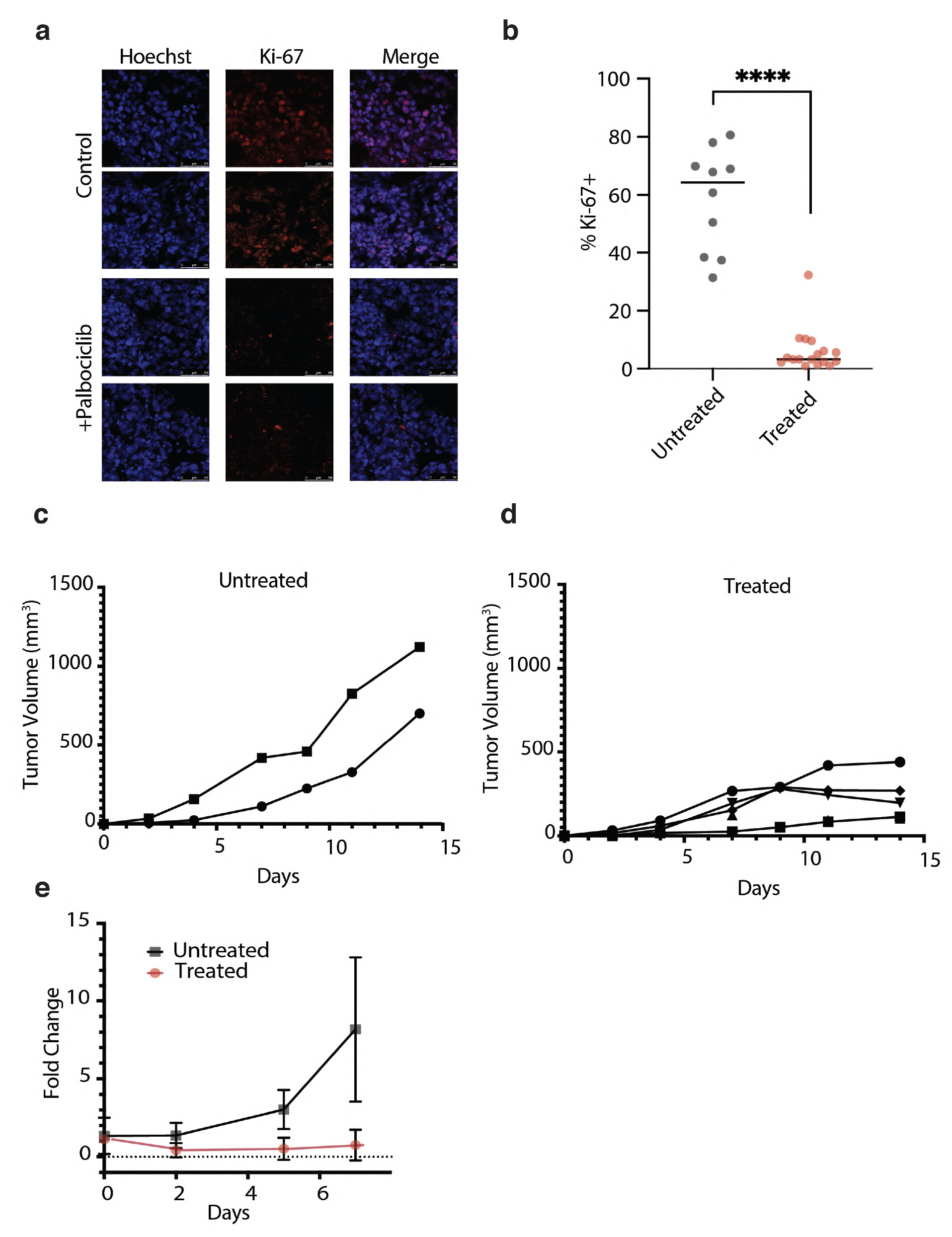
Characterization of tumor senescence using standard senescence markers. **(a)** Ki-67 staining with immunofluorescence. Treated tumors were compared to untreated and multiple slices and areas were used. (**b**) Quantification data obtained from confocal images (p<0.0001). Growth curves of the untreated and treated tumors indicating the proliferation of the untreated tumor and the growth arrest of treated (senescent) tumor (**c** and **d**). (**e**) Fold change representation of the data from b and c to highlight the growth arrest in treated tumors and compare it to the untreated tumors. Palbociclib treated (senescent) tumors show a marked decrease in proliferation rate indicative of senescence induction. Two tailed t test was used to calculate the significance (*p < .05, **p < .01, ***p < .001, and ****p<0.0001).

**Extended Data Figure 25.**
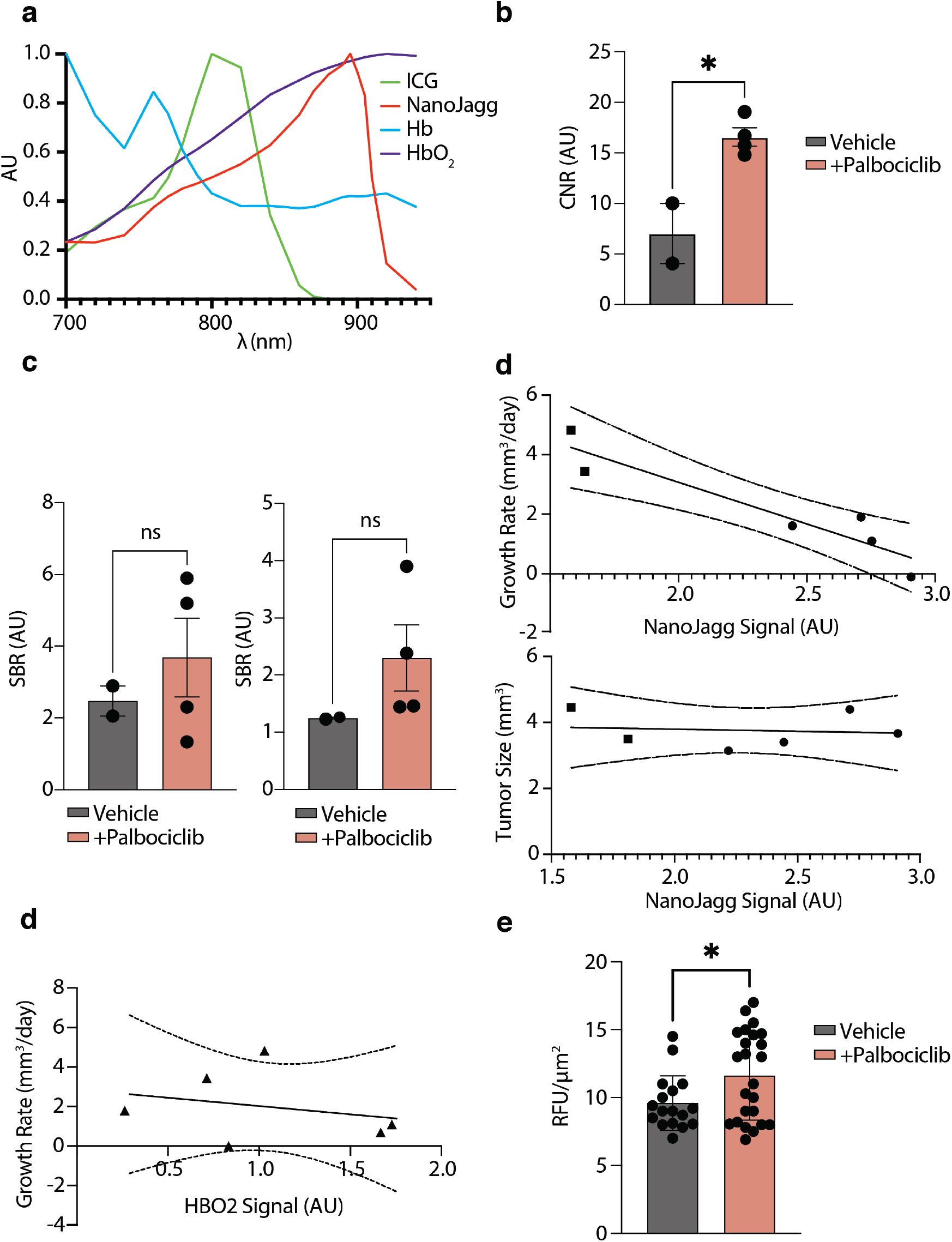
*In vivo* photoacoustic imaging using NanoJaggs. (**a**) Absorption spectra used for spectral unmixing employing existing reference spectra of ICG, Hb, and HbO2 and measured spectrum of the NanoJaggs. (**b**) Contrast to noise ratio (CNR) for untreated and treated (senescent) tumors indicating significantly higher CNR for treated tumors (p=0.0138). (**c**) The signal to background ratio (SBR) of linear spectral unmixed oxygenated hemoglobin (HbO2) and deoxygenated hemoglobin (Hb) are not significantly increased in treated tumors (p=0.5099 and p=0.2926). (**d**) NanoJagg signal does not correlate with tumor size (R^2^=0.015, p=0.7939, ns), but does correlate with the tumor growth rate (R^2^=0.8, p=0.002,**). (**e**) HBO2 signal does not correlate with tumor growth rate (R^2^=0.097, p=0.4952, ns) (**f**) Quantification of fluorescence per cell in untreated and treated tumors indicating higher fluorescence signal in each investigated cell of the treated tumors (p=0.0301). Data represent mean ± SD, and a Two tailed t test was used to calculate the significance (*p < .05, **p < .01, ***p < .001).

**Extended Data Figure 26.**
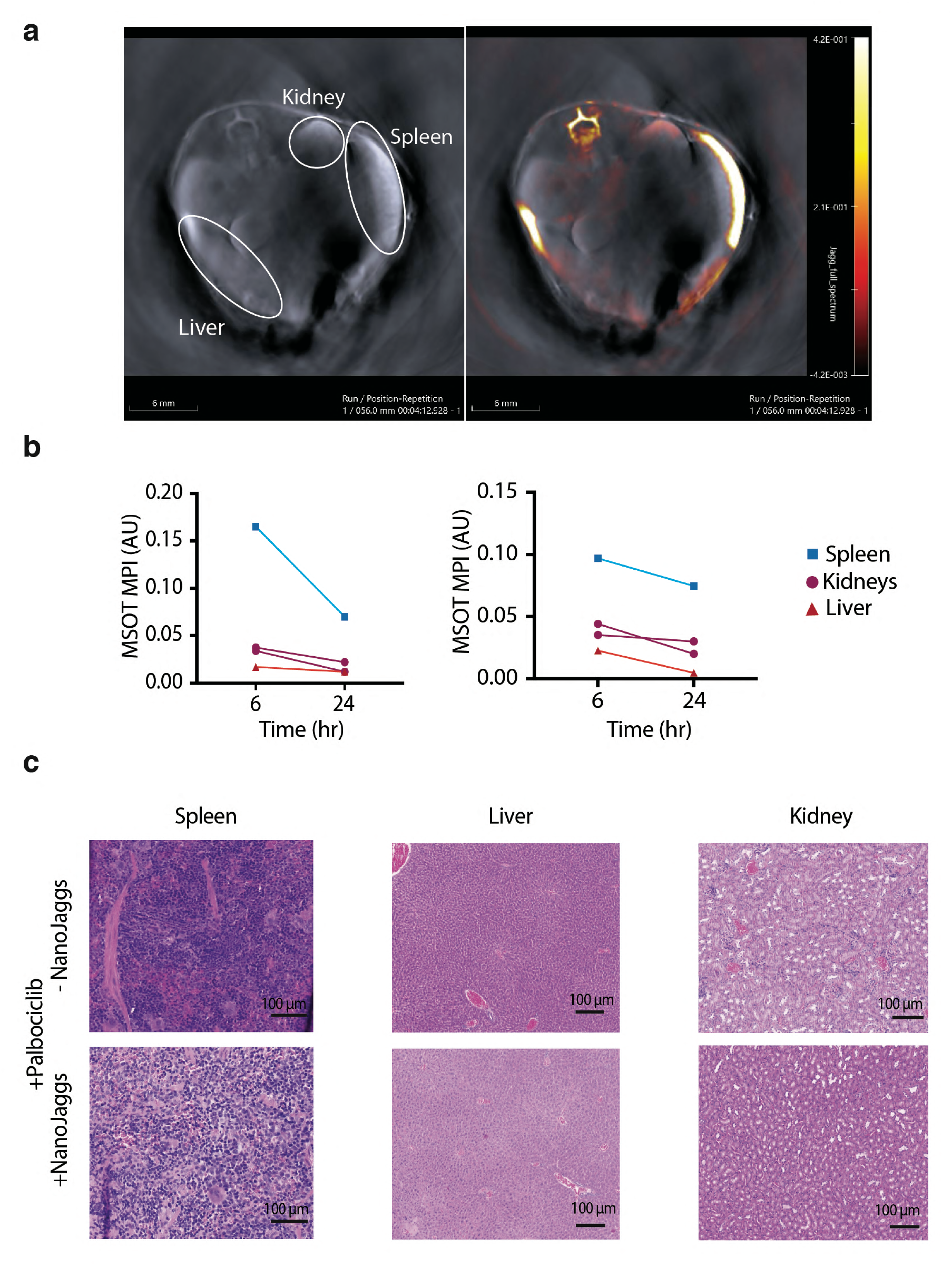
Evaluation of NanoJagg elimination and acute toxicity. (**a**) Spleen, liver and kidneys are involved in elimination of NanoJaggs as indicated by higher fluorescence in these organs. (**b**) NanoJagg signal in spleen, liver and kidneys drops significantly after 24 h in two studied mice. (**c**) H&E staining of spleen, liver and kidney in presence and absence of NanoJagg. NanoJagg treated organs do not appear structurally different indicating there is no acute toxicity for the length of the treatment.

